# A Sonification Framework for GPCR Molecular Dynamics: Auditory Signatures of β2-Adrenergic Receptor

**DOI:** 10.64898/2026.05.23.727422

**Authors:** Ekrem Yasar

## Abstract

Sonification, the systematic mapping of data to non-speech sound, has been applied with quantitative success in astronomy, seismology, and most recently materials chemistry, but has seen limited use in the analysis of biomolecular dynamics. Earlier protein-music studies have focused largely on the static amino-acid sequence, and G-protein-coupled receptor (GPCR) molecular dynamics (MD) trajectories have not previously been the subject of an auditory display framework. Here we present an end-to-end open-source sonification framework for GPCR molecular dynamics together with a quantitative cross-modal validation procedure, and we apply the framework as a proof of concept to three reference β2-adrenergic receptor (β2AR) trajectories from the GPCRMD repository spanning the activation continuum (inactive, active apo, active + orthosteric agonist). The framework extracts activation-related geometric features per MD frame, maps them under a single rule onto pitch, note duration, velocity, harmonic intensity, and percussive accents, and renders the result with three timbres (piano, violin, flute). The mapping was tested on two designed pairwise contrasts (activation pair; ligand pair) using Mann–Whitney U tests, Random Forest cross-modal classification with leave-one-instrument-out generalisation, and canonical correlation analysis between the MD and audio feature spaces. All four informative MD features differed between paired states at q < 1 × 10^−20^. A Random Forest classifier trained on audio features alone recovered the MD state with balanced accuracy 0.995 ± 0.003 (activation pair) and 1.000 ± 0.000 (ligand pair), corresponding to information-retention ratios of 1.006 and 1.000 relative to the MD-feature baseline. First canonical correlations between MD and audio spaces reached r_1_ = 0.926 (activation) and r_1_ = 0.995 (ligand). The sonification framework therefore provides a quantitatively faithful auditory representation of GPCR activation dynamics, with potential applications in exploratory MD analysis, accessibility, and education. The framework is system-agnostic and transfers to other GPCRs and allosteric MD systems without code changes beyond residue selection.

## 1. Introduction

Auditory display — the systematic mapping of data onto non-speech sound — has matured over three decades into a recognised modality for the perception, exploration, and communication of complex datasets [1-3]. The case for adding sound to a primarily-visual analysis pipeline rests on two complementary capacities of the auditory system: it parallel-processes temporal structure on timescales that vision serialises through animation, and it is sensitive to small relative changes in pitch, rhythm, and timbre that are difficult to encode within a single static visual frame [1]. These properties have supported useful sonification frameworks in fields as diverse as solar and gravitational-wave astronomy [4,5], seismology, climate time-series analysis, cellular calcium imaging, [6] and — more recently — chemistry and materials science, where mechanical-strain sonification of synthetic and biological materials has reached general-audience venues [7,8]. The common thread is that sound supplements rather than supplants visualisation: it offers a second perceptual channel along which patterns may emerge before they are evident in any single visual rendering [9].

Despite this maturity in adjacent fields, sonification remains relatively underused in the analysis of biomolecular dynamics. Molecular dynamics (MD) simulations now routinely produce microsecond-to-millisecond conformational trajectories for systems of biological interest; [10,11] the standard output remains a cartoon or surface animation interpreted by visual inspection alone. The opportunity is clear: an MD trajectory is, by construction, a long temporal sequence of structural states — exactly the kind of data for which auditory display has historically offered notable perceptual benefits.

The existing body of biomolecular sonification work has focused largely on primary structure — the linear amino-acid sequence — rather than on conformational dynamics. Pioneering efforts mapped the sequence directly to pitch using either physico-chemical or arbitrary lookup tables; [12,13] more recent work has used machine-learning representations of sequence to drive musical composition [7,8]. These studies have established that proteins can be rendered into music in aesthetically interesting ways, but they share two limitations relative to what the present work attempts.

First, they sonify a static representation of a protein (the sequence, or a single reference structure) rather than a dynamical one. The conformational ensemble — the central output of MD and the substrate from which functional events such as receptor activation are read — has not, to our knowledge, been the target of a per-residue or per-frame sonification framework [14]. Second, prior sonifications have been evaluated primarily on aesthetic or qualitative grounds; quantitative validation of biomolecular sonification as an information-preserving representation, in the sense that downstream computational analyses on the rendered audio recover the latent state present in the source data, has not been a standard part of the evaluation.

A particularly underserved system in this landscape is the G-protein-coupled receptor (GPCR) family. GPCRs are the largest class of membrane receptors and drug targets, [15] their activation has been mapped at near-atomic resolution by both crystallographic and cryo-EM ensembles, [16-18] and large-scale MD trajectories of multiple receptor states are publicly available through the GPCRMD repository [19]. Despite this, GPCR molecular dynamics has not, to our knowledge, previously been explored through an auditory display framework. The present work was motivated by the observation that GPCRs are simultaneously (i) a canonical model system for studying receptor activation through MD, and (ii) absent from the auditory display literature.

We selected the β2-adrenergic receptor (β2AR) as the test system because it is one of the best-characterised GPCR activation models [20-22]. β2AR has structurally resolved inactive, active apo, and active agonist-bound states; its activation-related geometric markers — TM3–TM6 intracellular distance, the DRY-motif ionic lock between residues R3.50 and E6.30, the conserved NPxxY motif of TM7, and the orthosteric ligand binding pose — are quantitatively validated and individually interpretable [23]. GPCRMD provides three high-quality reference trajectories spanning the activation landscape (Dynamic IDs 11, 116, 117), allowing comparison of three well-defined states under matched simulation conditions.

The β2AR activation cascade is also pedagogically clean: TM6 swings outward at its intracellular end, the ionic lock disengages, the NPxxY motif rearranges to a G-protein-binding-competent rotamer, and — when an orthosteric agonist is bound — these conformational changes are stabilised and the receptor commits to the active ensemble. This is precisely the kind of multi-feature, monotonically progressing conformational transition for which a multi-axis sonification mapping (pitch, tempo, loudness, harmonic intensity, percussion) can encode each feature on a distinct perceptual axis. β2AR therefore offers a stringent test of whether such a mapping can simultaneously preserve the information content of multiple activation-related observables, while remaining a sufficiently familiar system that biological interpretations of the rendered audio can be readily cross-validated against existing literature.

Here we present a sonification framework for GPCR molecular dynamics designed to be system-agnostic — transferable to any GPCR (or any allosteric MD system) by adjusting only the residue-selection strings — and we test the framework on β2AR as a proof of concept. We hypothesised that activation-related geometric features extracted from GPCR MD trajectories, when mapped under a fixed rule onto musical parameters and rendered into audio, produce a representation that preserves the underlying MD-state information in a quantifiable, instrument-independent manner. We assessed this hypothesis on the three GPCRMD β2AR trajectories using two designed pairwise contrasts (activation pair: inactive vs active apo; ligand pair: active apo vs active + orthosteric agonist), rendered under an identical mapping with three timbres (piano, violin, flute) to dissociate MD signal from synthesis-specific artefacts.

The contributions of this work are fourfold: (i) an end-to-end open-source sonification framework for GPCR molecular dynamics, distributed as a set of reproducible Google Colab notebooks together with a synchronised HTML player, designed to be applied directly to any GPCR by adjusting only the residue-selection strings; (ii) a contrastive per-pair experimental design that tests information preservation for the two scientifically meaningful comparisons directly, rather than only a global multi-class problem; (iii) a cross-instrument robustness analysis that separates MD signal from instrument-specific timbre artefacts; and (iv) a quantitative cross-modal validation procedure — combining Random Forest classification, leave-one-instrument-out generalisation, and canonical correlation analysis between MD and audio feature spaces — providing interpretation-free metrics for evaluating biomolecular sonification mappings. The β2AR demonstration in this manuscript is intended as a proof of concept for the framework; the pipeline transfers to any GPCR or any allosteric MD system without code changes beyond residue selection.

## 2. Methods

### 2.1. Trajectory data

The three molecular dynamics trajectories of the β2-adrenergic receptor analysed in this work were retrieved from the GPCRMD repository: [19] the inactive apo state (GPCRMD Dynamic ID 11), the active apo state (Dynamic ID 116), and the active state with the orthosteric agonist bound (Dynamic ID 117; ligand residue name “ALE”). Each trajectory comprises 2 500 frames simulated under matched conditions (lipid bilayer, explicit-solvent, 310 K) by the GPCRMD consortium; full simulation provenance is provided in the original repository entries. Trajectories were used at a uniform stride of 5, yielding 500 analysis frames per state and a total of 1 500 frames across the three states. We refer to these throughout as the inactive, active, and active + ligand states.

### 2.2. MD feature extraction

Each trajectory was loaded with MDAnalysis 2.8 and rigid-body-aligned in memory on all Cα atoms using MDAnalysis.analysis.align.AlignTraj with the reference taken as the first frame of the same trajectory. In-memory alignment removes whole-receptor translation and tumbling so that subsequent RMSDs reflect internal conformational change rather than rigid-body drift.

Six per-frame features were computed: (i) global Cα RMSD with respect to the reference frame, computed without further superposition; (ii) TM3–TM6 intracellular distance, defined as the Cα geometric-centre distance between residues 130–136 (the TM3 cytoplasmic end including the DRY motif) and residues 262–272 (the TM6 cytoplasmic end including E6.30 and L6.34); (iii) NPxxY motif RMSD over residues 322–326 (N322-P323-L324-I325-Y326), computed against the reference frame after global alignment; (iv) DRY ionic-lock distance, the Cα–Cα distance between R3.50 (Arg131) and E6.30 (Glu268); (v) ligand minimum heavy-atom distance to receptor heavy atoms (scipy.spatial.distance.cdist minimum); and (vi) ligand–receptor contact count, defined as the number of receptor heavy atoms within 4.5 Å of any ligand heavy atom. Features (v) and (vi) are populated only for the ligand-bound trajectory. Residue-range selections were verified by direct inspection of each topology PDB and confirmed identical across the three GPCRMD trajectories used.

### 2.3. Sonification mapping

Each per-frame feature was mapped onto a musical parameter under a single deterministic rule applied identically to every state. Continuous features were first globally min–max normalised against the union of all three states; identical MD values therefore map to identical musical values across states, and any difference between rendered states is attributable to the MD rather than to per-state normalisation. The mapping is: TM3–TM6 distance → pitch, quantised to the C-major pentatonic scale within the MIDI range 48−84 (C3–C6); NPxxY RMSD → note duration, linearly scaled to 0.12−0.45 s; Cα RMSD → note velocity, linearly scaled to MIDI 40–110; DRY ionic-lock distance → harmonic-pad intensity (8-note moving window; New-Age-Pad GM patch 89); Ligand contact count → percussive accents (acoustic snare, GM drum note 38), triggered when the per-frame count is non-zero. Inter-note gap was fixed at 0.02 s; tempo at 90 BPM. Three timbres were rendered from the same mapping table using the General-MIDI program changes 0 (piano), 40 (solo violin) and 73 (flute). MIDI scores were generated with pretty_midi 0.2.10^27^ and synthesised to 44.1 kHz WAV with FluidSynth 2.3 using the FluidR3 GM SoundFont.

### 2.4. Acoustic feature extraction

Each rendered WAV (9 single-state files in total) was sliced into 1.0-s analysis windows with 0.5-s hop, yielding approximately 230–340 windows per (state × instrument) and ∼1 700 paired windows per pair. For each window, a 28-dimensional descriptor vector was computed with librosa 0.10:^28^ 13 mel-frequency cepstral coefficients reported as per-window mean and standard deviation (26 features), spectral centroid (mean, σ), spectral rolloff at 85 % (mean, σ), spectral bandwidth (mean, σ), root-mean-square energy (mean, σ), and zero-crossing rate (mean, σ).

### 2.5. Statistical analysis

Two pairwise contrasts were tested in parallel: the activation pair (inactive vs active apo) and the ligand pair (active apo vs active + ligand). For each pair × each feature (MD or audio), a two-sided Mann–Whitney U test was performed (scipy.stats.mannwhitneyu), accompanied by Cliff’s δ as a non-parametric effect size.^29^ p-values were combined per test family and adjusted by the Benjamini–Hochberg false-discovery-rate procedure^30^ to a per-family threshold of q < 0.05.

For the MD-side analysis, all four activation markers (Cα RMSD, TM3–TM6 distance, NPxxY RMSD, DRY ionic lock) carry information in every pair and were tested in both. The two ligand-specific features (Ligand_min_distance, Ligand_contact_count) are structurally undefined in apo systems and were excluded from the activation-pair tests; for the ligand pair they were included with the apo state populated by biologically motivated sentinel values described in §2.6.

### 2.6. Cross-modal classification, LOIO generalisation, and canonical correlation

The central computational test of the hypothesis is whether a classifier trained on audio features alone can recover the MD state. For each pair we trained a Random Forest classifier (sklearn.ensemble.RandomForestClassifier, n_estimators = 300, random_state = 42) with stratified 5-fold cross-validation, scored by balanced accuracy.^31^ Models were fitted four times per pair: once per instrument (piano, violin, flute) and once pooled across all three instruments.

To establish an information-theoretic upper bound for the audio classifier, we trained the same Random Forest on the raw MD features for each pair. The set of MD features available to the MD baseline is selected automatically per pair: a feature is included only if at least one of the pair’s two states has non-missing observations. For the activation pair, this rule retains the four activation markers and drops the two ligand-specific features (apo states have no ligand). For the ligand pair, all six features are retained, with apo-state ligand values imputed by biological sentinels: Ligand_contact_count = 0 (literal absence of ligand atoms) and Ligand_min_distance = 30 Å (an arbitrarily large “no-ligand” sentinel, chosen well above any physical bound distance). Median-within-state imputation was used for any other residual missingness. The audio-pooled balanced accuracy divided by the per-pair MD-baseline balanced accuracy defines an information-retention ratio per pair; values near 1.0 indicate that the sonification mapping preserves the MD’s binary discriminative information.

To probe whether the audio-state signal is instrument-independent, we ran leave-one-instrument-out (LOIO) cross-validation per pair: a Random Forest was trained on two of the three instruments’ windows and tested on the held-out third (three LOIO models per pair, six per study).

Finally, canonical correlation analysis (sklearn.cross_decomposition.CCA, n_components = 4, max_iter = 2000) was used to quantify the bilateral correspondence between the MD and audio feature spaces in classifier-free terms. Paired (MD-frame, audio-window) samples were constructed by aligning each audio window’s centre time to the nearest MIDI-note start time reconstructed from the per-state mapping table (using start[i] = Σ□< □ duration[j] + i · gap with gap = 0.02 s). Standardised MD and audio matrices were fitted with CCA per pair and for the three-state union; the first four canonical correlations are reported.

### 2.7. Code and data availability

All analyses are distributed as five reproducible Google Colab notebooks (01_GPCRMD_feature_extraction.ipynb through 05_GPCR_sonification_classification.ipynb) together with a synchronised HTML player and pre-rendered audio/video assets. The repository is archived at Zenodo 10.5281/zenodo.20343439, and source code is mirrored on GitHub at https://github.com/eygpcr/gpcr-sonification-beta2ar. The directory layout follows the convention data/raw/{state}/, data/processed/, outputs/audio/, outputs/figures/, outputs/tables/, and outputs/video/. All figure-generating code uses an identical publication-style matplotlib configuration (sans-serif, 9–10 pt, embedded Type 42 fonts, 600 dpi PNG + PDF + SVG); the Okabe–Ito colour-blind-safe palette is used throughout for state colours and a complementary categorical palette for pair-level overlays.

## 3. Results

### 3.1. MD features separate the three β2AR states with strong, monotonic effect sizes

We first verified that the six geometric features extracted from the three GPCRMD β2AR trajectories (N = 500 sonification-strided frames per state) carry the activation-state information they are intended to encode. For both designed contrasts — the activation pair (inactive vs active apo) and the ligand pair (active apo vs active + orthosteric agonist) — we performed two-sided Mann–Whitney U tests on every feature, accompanied by Cliff’s δ as a non-parametric effect size, with Benjamini–Hochberg correction within each test family (Table 3).

**Table 1.**
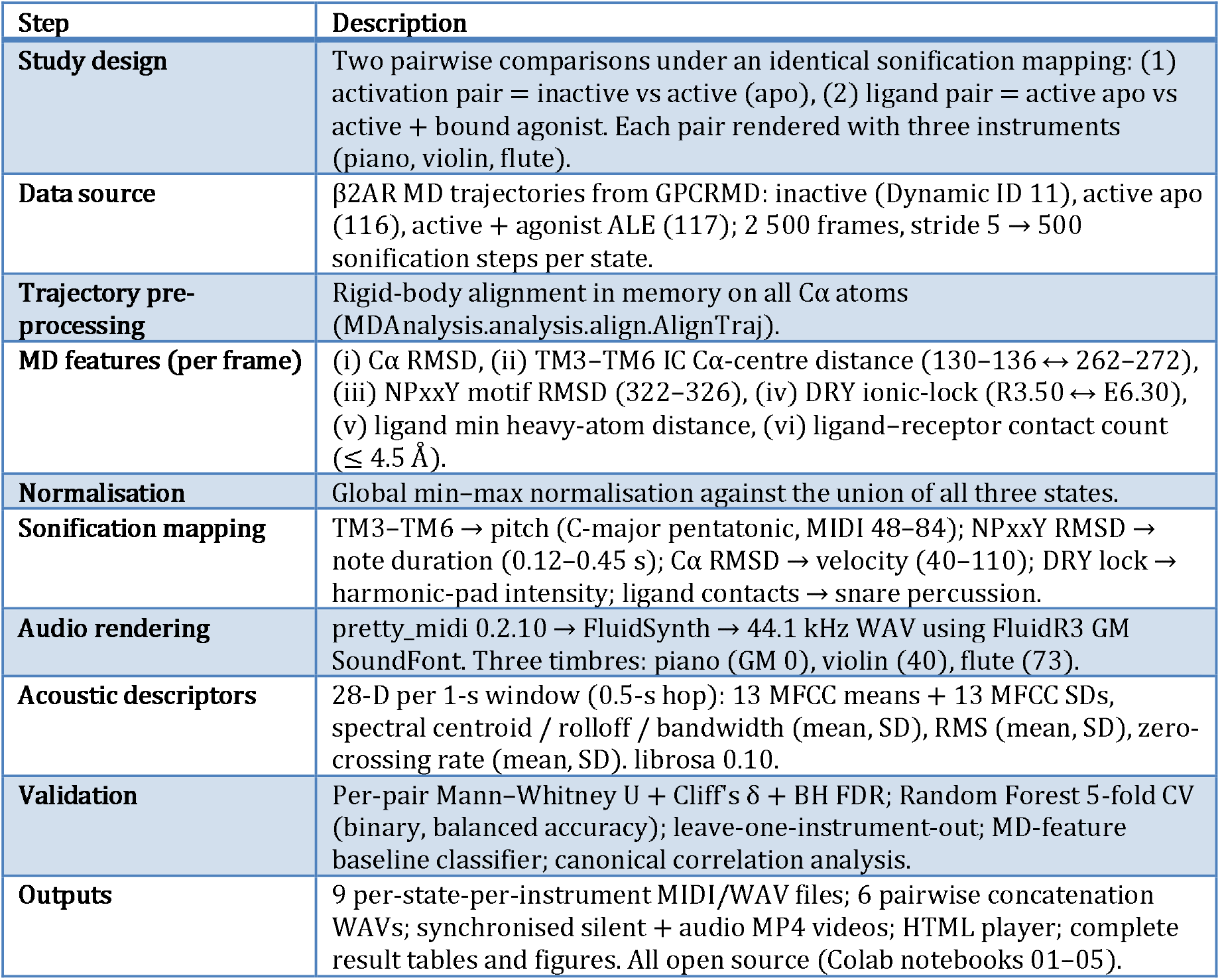
Study design and pipeline summary. A consolidated overview of the experimental design, data, computational pipeline, and output deliverables.

| Step | Description |
| --- | --- |
| **Study design** | Two pairwise comparisons under an identical sonification mapping: (1) activation pair = inactive vs active (apo), (2) ligand pair = active apo vs active + bound agonist. Each pair rendered with three instruments (piano, violin, flute). |
| **Data source** | β2AR MD trajectories from GPCRMD: inactive (Dynamic ID 11), active apo (116), active + agonist ALE (117); 2 500 frames, stride 5 → 500 sonification steps per state. |
| **Trajectory pre-processing** | Rigid-body alignment in memory on all Cα atoms (MDAnalysis.analysis.align.AlignTraj). |
| **MD features (per frame)** | (i) Cα RMSD, (ii) TM3–TM6 IC Cα-centre distance (130–136 ↔ 262–272), (iii) NPxxY motif RMSD (322–326), (iv) DRY ionic-lock (R3.50 ↔ E6.30), (v) ligand min heavy-atom distance, (vi) ligand–receptor contact count ($\leq 4.5$ Å). |
| **Normalisation** | Global min–max normalisation against the union of all three states. |
| **Sonification mapping** | TM3–TM6 → pitch (C-major pentatonic, MIDI 48–84); NPxxY RMSD → note duration (0.12–0.45 s); Cα RMSD → velocity (40–110); DRY lock → harmonic-pad intensity; ligand contacts → snare percussion. |
| **Audio rendering** | pretty_midi 0.2.10 → FluidSynth → 44.1 kHz WAV using FluidR3 GM SoundFont. Three timbres: piano (GM 0), violin (40), flute (73). |
| **Acoustic descriptors** | 28-D per 1-s window (0.5-s hop): 13 MFCC means + 13 MFCC SDs, spectral centroid / rolloff / bandwidth (mean, SD), RMS (mean, SD), zero-crossing rate (mean, SD). librosa 0.10. |
| **Validation** | Per-pair Mann–Whitney U + Cliff's δ + BH FDR; Random Forest 5-fold CV (binary, balanced accuracy); leave-one-instrument-out; MD-feature baseline classifier; canonical correlation analysis. |
| **Outputs** | 9 per-state-per-instrument MIDI/WAV files; 6 pairwise concatenation WAVs; synchronised silent + audio MP4 videos; HTML player; complete result tables and figures. All open source (Colab notebooks 01–05). |

**Table 2.** Per-state means and standard deviations of MD features. Distances and RMSDs in Å; ligand contact count as integer atom count. Each cell: mean ± SD over 500 strided MD frames. Dashes indicate features that are structurally undefined for that state (no ligand in apo systems). The NPxxY RMSD inversion in the ligand-bound state (4.08 → 2.08 Å) is consistent with agonist-induced motif rigidification (see Discussion §4.2).

| MD feature | Inactive | Active (apo) | Active + ligand |
| --- | --- | --- | --- |
| **Cα RMSD (Å)** | 3.44 ± 0.78 | 4.06 ± 0.90 | 4.80 ± 1.13 |
| **TM3–TM6 intracellular distance (Å)** | 11.86 ± 1.30 | 13.80 ± 1.54 | 16.17 ± 1.95 |
| **NPxxY motif RMSD (Å)** | 1.96 ± 0.79 | 4.08 ± 1.45 | 2.08 ± 0.68 |
| **DRY ionic-lock distance R3.50 → E6.30 (Å)** | 8.49 ± 1.22 | 10.19 ± 1.59 | 13.05 ± 1.69 |
| **Ligand minimum distance to** | — | — | 1.81 ± 0.17 |

| receptor (Å) |  |  |  |
| --- | --- | --- | --- |
| Ligand–receptor contact count<br>(atoms $\leq 4.5$ Å) | — | — | 188.5 ± 29.1 |

**Table 3.**
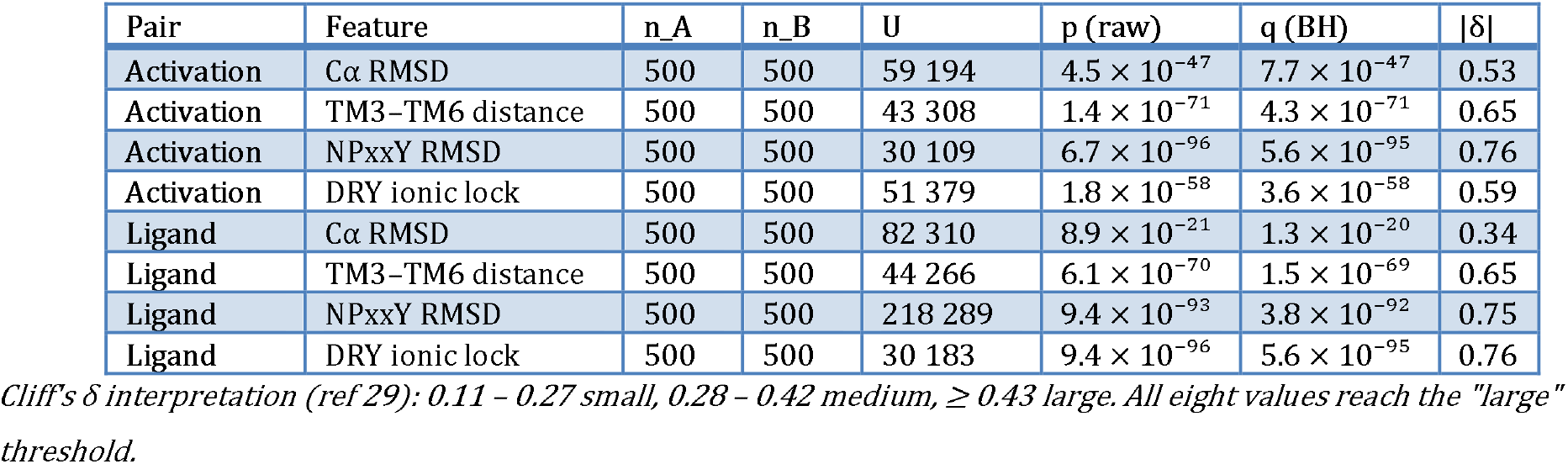
Per-pair Mann–Whitney U tests of MD features. Two-sided MWU on each MD feature per pair, with Cliff’s δ as non-parametric effect size; p-values adjusted within each test family by the Benjamini–Hochberg FDR procedure. All four informative features differed significantly in both pair contrasts at q < 1 × 10^−20^. Ligand-specific features (Ligand_min_distance, Ligand_contact_count) are structurally determined for the ligand pair and are reported in Table 2; they are excluded from this MWU table because at least one state lacks values structurally.

For the activation pair, all four activation markers separated the two states at q < 1 × 10^−46^: TM3–TM6 intracellular distance increased from 11.86 ± 1.30 Å in the inactive state to 13.80 ± 1.54 Å in the active state (| δ| = 0.65), the DRY ionic-lock distance opened from 8.49 ± 1.22 Å to 10.19 ± 1.59 Å (| δ| = 0.59), the NPxxY-motif RMSD increased from 1.96 ± 0.79 Å to 4.08 ± 1.45 Å (| δ| = 0.76, the largest single-feature effect), and the global Cα RMSD rose from 3.44 ± 0.78 Å to 4.06 ± 0.90 Å (| δ| = 0.53). All four effect sizes fall in the medium-to-large range.^29^

For the ligand pair, the same four features remained significant at q < 1 × 10^−20^, with TM3– TM6 widening further to 16.17 ± 1.95 Å (| δ| = 0.65) and the ionic lock disengaging almost fully to 13.05 ± 1.69 Å (| δ| = 0.76). A biologically informative inversion emerged in the NPxxY motif: while the apo active state shows the highest NPxxY RMSD (4.08 ± 1.45 Å), agonist binding stabilises this motif (RMSD 2.08 ± 0.68 Å, | δ| = 0.75). This is consistent with prior reports that orthosteric agonists rigidify the TM7 NPxxY rotamer into a G-protein-binding-competent conformation^23^,^24^ and emerges directly from our per-frame quantification (Figure 2). The two ligand-specific features are, by construction, only populated for the bound trajectory and show the expected stable orthosteric engagement (1.81 ± 0.17 Å, 188 ± 29 atoms within 4.5 Å). Together, these results establish that the MD feature panel encodes a monotonic three-state activation continuum (inactive → active → active + ligand) that is quantitatively recoverable, providing a solid substrate for the downstream sonification mapping.

**Figure 2.**
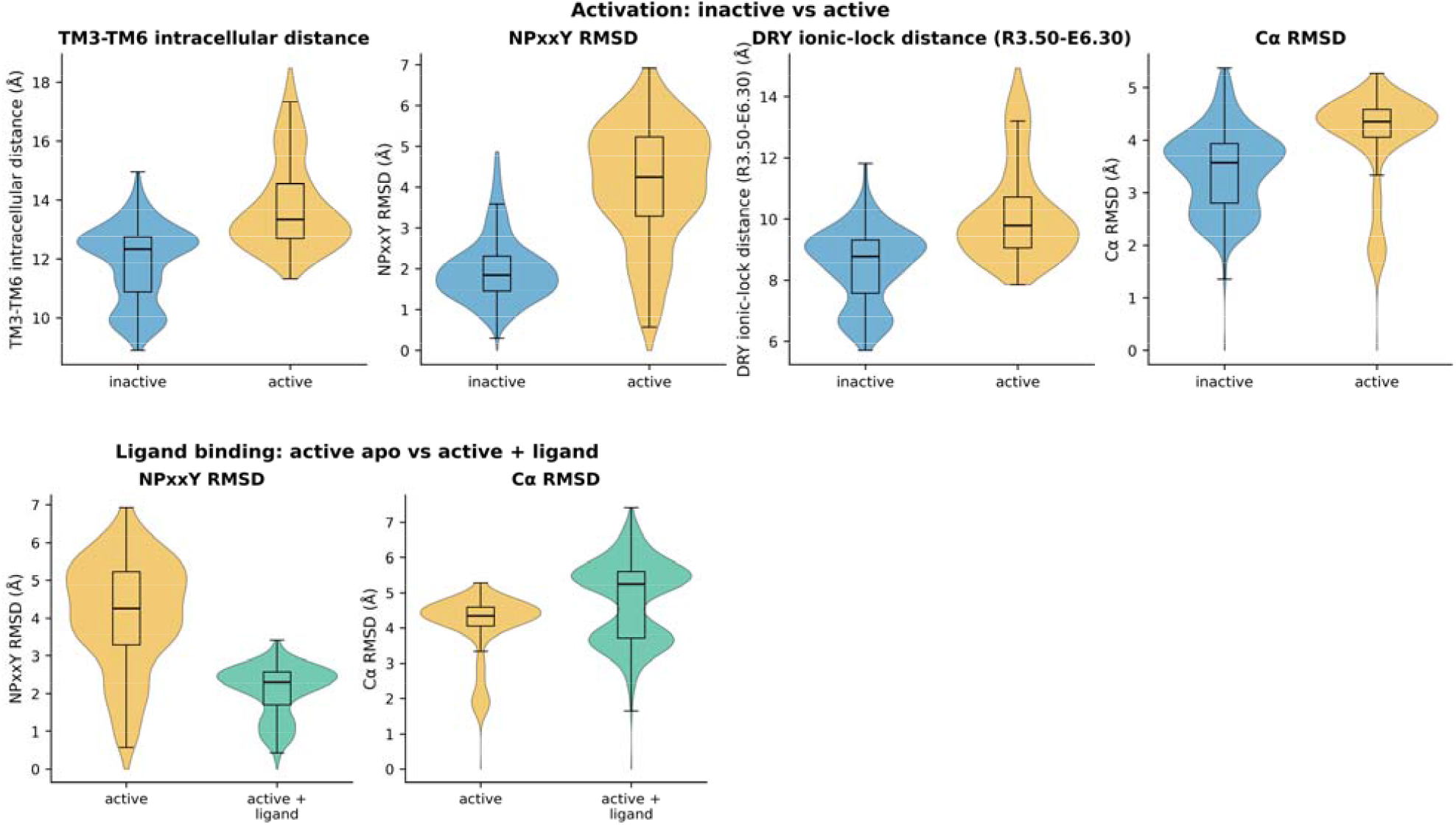
MD-feature distributions per pair. Distributions of the four informative MD features that separate the two states in each designed pair. (A) Activation pair (inactive vs. active apo): TM3–TM6 intracellular distance, NPxxY motif RMSD, DRY ionic-lock distance (R3.50 ->E6.30), and Cα RMSD. (B) Ligand pair (active apo vs. active + orthosteric agonist): same four features. Each panel shows violin plots overlaid with box plots (median, IQR, 1.5×IQR whiskers, fliers suppressed) computed over 500 strided MD frames per state. Note in panel B that NPxxY RMSD decreases on ligand binding (4.08 → 2.08 Å, |Cliff’s δ| = 0.75) — opposite in direction to the other three markers — consistent with agonist-induced rigidification of the TM7 NPxxY rotamer. State colours (Okabe–Ito): inactive (blue), active apo (orange), active + ligand (green). All four features differed significantly between states at q < 1 × 10^−20^ within each pair (two-sided Mann–Whitney U, Benjamini–Hochberg FDR; see Table 3).

### 3.2. The sonification mapping produces state-specific audio signatures

We next asked whether the deterministic mapping from MD features to musical parameters (Methods 2.3) translates the MD-state separation of §3.1 into the rendered audio. Under the identical mapping rule applied to every state, the three MIDI score families differ along three orthogonal musical axes simultaneously (Table S1): mean pitch progresses from 57.7 (A3) in the inactive state through 64.0 (E4) in the active state to 71.7 (B4) in the active + ligand state; mean note duration is shortest in the inactive (0.21 s), longest in the active (0.31 s; reflecting the NPxxY-driven dynamics maximum), and short again in the active + ligand state (0.22 s, consistent with the NPxxY stabilisation noted above); mean velocity rises monotonically from 72 to 78 to 85. Each state therefore occupies a distinct perceptual region of the (pitch × tempo × loudness) space.

The mapping rule is rendered through three timbres (piano, violin, flute) using General MIDI program changes only, so any per-state difference in the audio is attributable to the MD rather than to instrument-specific synthesis (Figure 3, Figure 4). We quantified the audio-side state separation by extracting 28 spectro-temporal descriptors (MFCCs, spectral centroid / rolloff / bandwidth, RMS, zero-crossing rate) from 1-s analysis windows with 0.5-s hop (∼600 windows per state per instrument, ∼1 700 paired windows per pair). MWU tests with BH correction were applied per pair × per descriptor.

**Figure 3.**
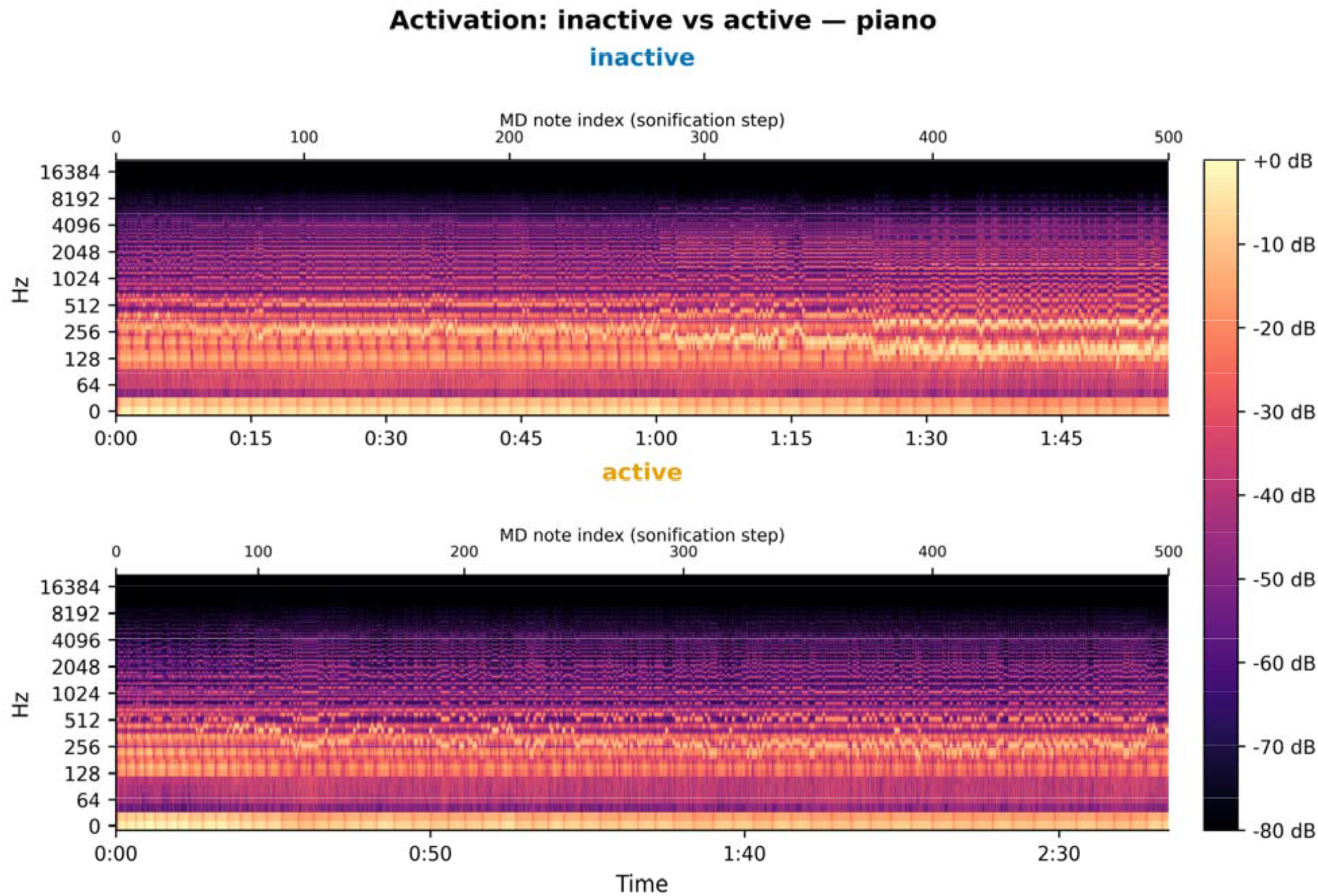
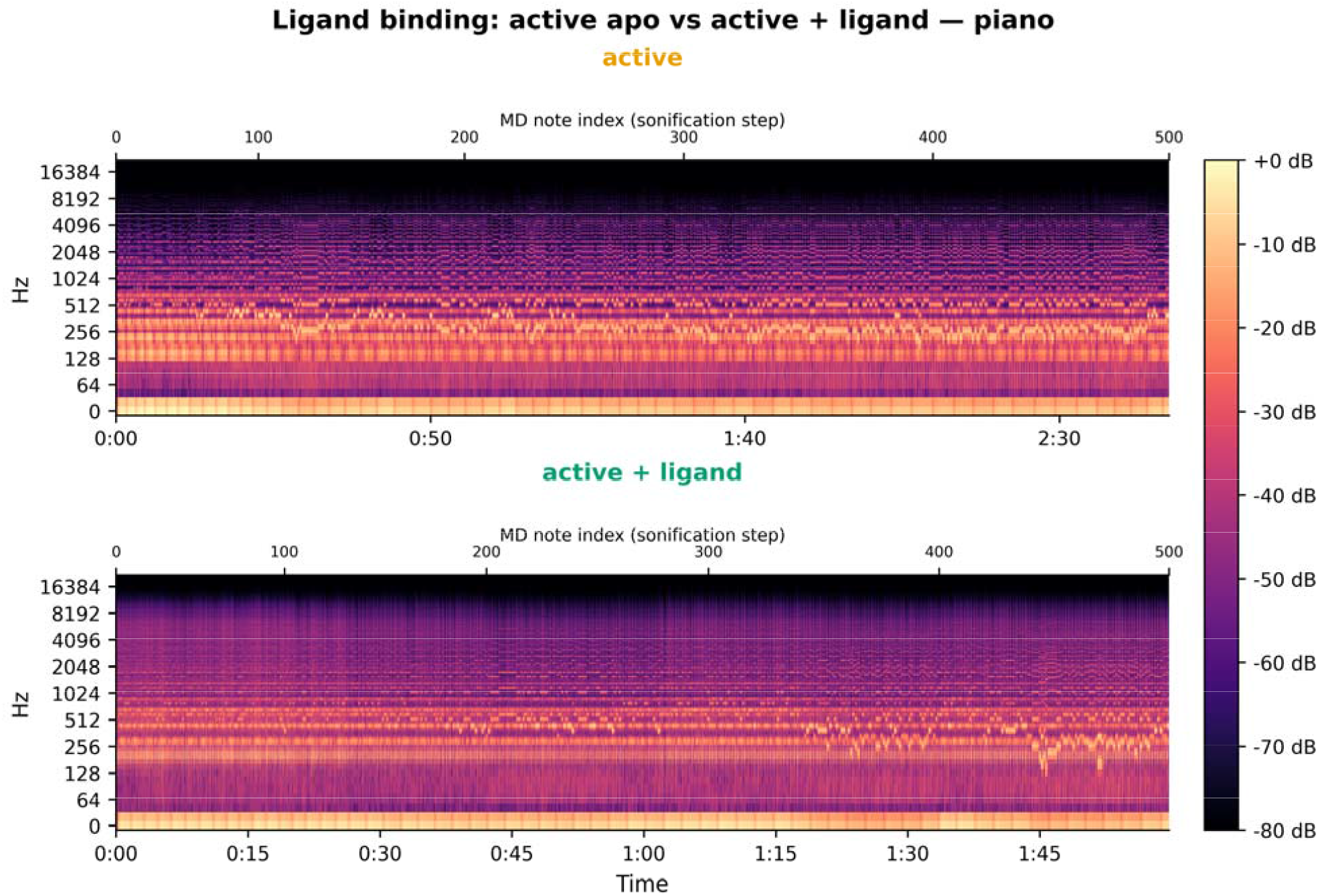
Side-by-side piano spectrograms per pair. Spectrograms of the rendered piano audio for the two designed pairwise contrasts under the identical sonification mapping (Methods §2.3). **(A) Activation pair:** inactive (top) vs. active apo (bottom). **(B) Ligand pair:** active apo (top) vs. active + orthosteric agonist (bottom). Each spectrogram was computed as the short-time Fourier transform (n_fft = 2048, hop = 512) of the FluidSynth-rendered WAV and converted to dB (librosa.amplitude_to_db, ref = max). Lower (bottom) x-axis: real audio time (s); upper (top) x-axis: corresponding MD note index (0–500), reconstructed from the per-state mapping table so that the same MD step lies at the same top-axis position across panels even though the two states have different total audio durations. State titles are tinted in the Okabe–Ito state colour (blue, orange, green). Note in (B) the dense, broadband percussive bursts in the lower panel — the snare track that is triggered exclusively by the ligand-contact feature.

**Figure 4.**
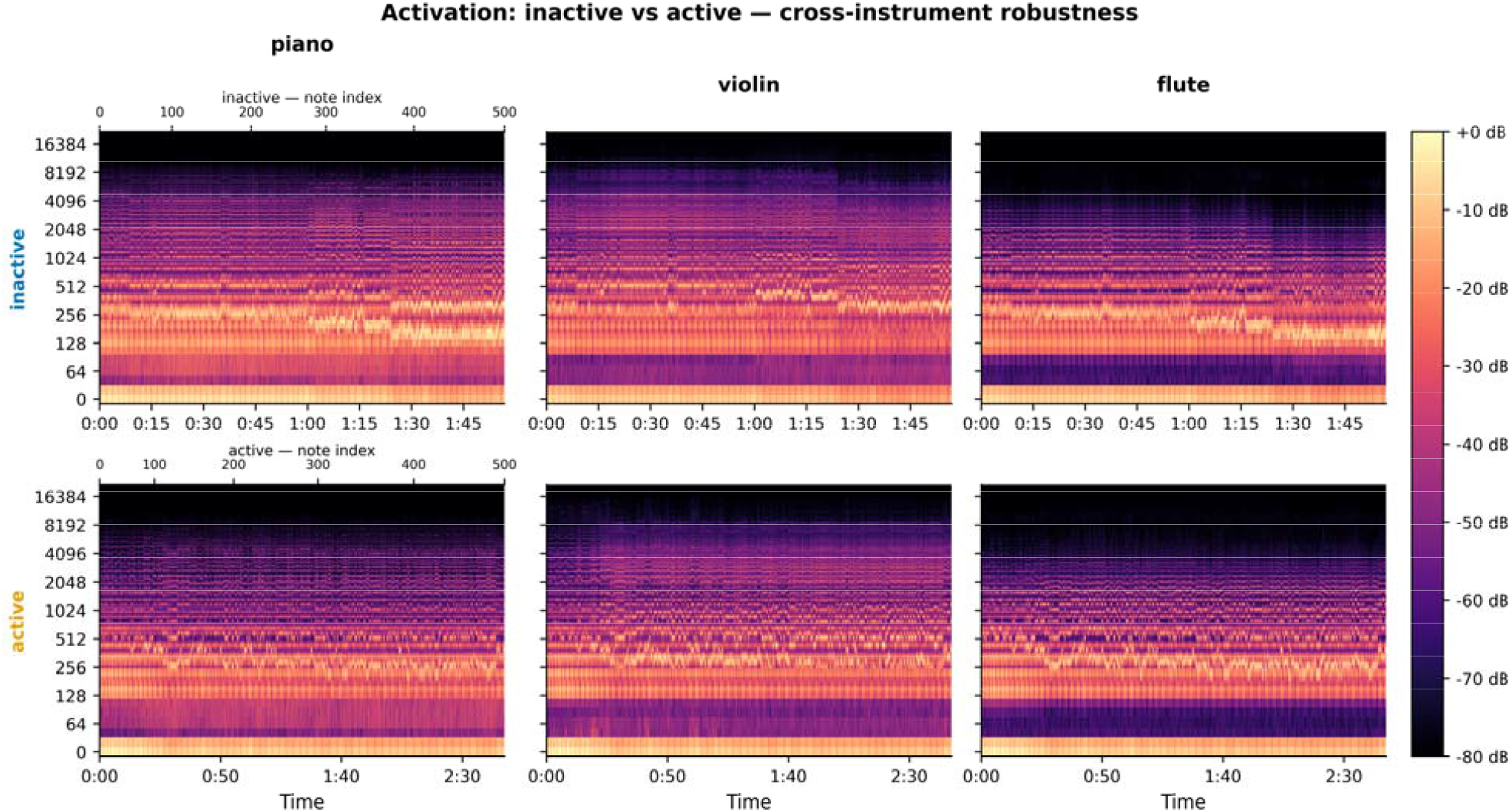
Cross-instrument robustness grid — activation pair. The activation-pair contrast (inactive vs. active apo) rendered with three timbres (piano, violin, flute) under the identical sonification mapping. Rows: state (inactive, active apo). Columns: instrument. Each cell shows the same spectrogram representation as Figure 3 with a shared log-frequency axis and a per-row shared time axis (state-specific audio durations). The within-column contrast between rows — visibly preserved across all three instruments — is the qualitative evidence that the auditory state distinction is carried by the MD-derived musical content rather than by instrument-specific timbre. Shared colour map (magma) and intensity scale across the grid. Row labels in Okabe–Ito state colour.

For both pairs, 34 of the 36 descriptors tested per pair reached q < 0.05 (Table S3). The pair separations differ in character: the ligand pair shows extremely strong effects (17 descriptors with | δ| > 0.7, including mfcc1_mean at p = 1.8 × 10^−273^ and | δ| = 1.00, spec_rolloff_std | δ| = 0.99, rms_mean | δ| = 0.99), dominated by the snare-percussion track that is triggered only when the ligand-contact feature is non-zero. The activation pair shows a more diffuse but consistent signature (1 large + 12 medium-effect descriptors, the largest being rms_mean with | δ| = 0.82), driven by the joint pitch–tempo–velocity shift. Both signature classes are visible in the side-by-side spectrograms of Figure 3 and persist across timbres in the cross-instrument grid of Figure 4.

### 3.3. An audio-only classifier recovers the MD state with near-perfect accuracy

The central computational test of our hypothesis is whether a classifier trained on audio descriptors alone — without access to the MD features themselves — can recover the MD state. For each pair we trained Random Forest classifiers (300 trees, fixed seed) under stratified 5-fold cross-validation, scored by balanced accuracy. Models were fitted per instrument (piano, violin, flute) and pooled across all three instruments (Table 4, Figure 5). For the activation pair, balanced accuracies were 0.988 ± 0.008 (piano), 0.996 ± 0.009 (violin), 0.998 ± 0.003 (flute), and 0.9954 ± 0.0026 (pooled across instruments, N = 1 695 windows). For the ligand pair, every single classifier — per-instrument and pooled — reached balanced accuracy 1.000 ± 0.000 (perfect recovery on N = 1 713 windows). The pooled confusion matrices (Figure 5) show 691/696 inactive windows and 997/999 active windows correctly assigned for the activation pair, and all 999 active and 714 active + ligand windows correctly assigned for the ligand pair.

**Table 4.** Cross-modal classification of MD state from audio features. Random Forest classifier (n_estimators = 300, random_state = 42) trained on 28-D windowed audio descriptors (1-s windows, 0.5-s hop) to recover the MD state. Per-pair binary classification (chance = 0.500); 5-fold stratified cross-validation; balanced accuracy reported as mean ± SD across folds. Per-instrument and pooled-across-instruments models for each pair, against the MD-feature baseline (same RF on raw MD features, per-pair valid feature subset with biological sentinel imputation for apo ligand features). LOIO = leave-one-instrument-out. Information-retention ratio = audio_pooled_acc / MD_baseline_acc.

**Panel A. Within-instrument and pooled (5-fold CV)**
| Pair | Model | n_samples | Balanced accuracy |
| --- | --- | --- | --- |
| Activation | audio / piano | 565 | 0.988 ± 0.008 |
| Activation | audio / violin | 565 | 0.996 ± 0.009 |
| Activation | audio / flute | 565 | 0.998 ± 0.003 |
| Activation | audio / pooled | 1 695 | 0.995 ± 0.003 |
| Activation | MD baseline | 1 000 | 0.989 ± 0.006 |
| Activation | Information-retention ratio | — | 1.006 |
| Ligand | audio / piano | 571 | 1.000 ± 0.000 |
| Ligand | audio / violin | 571 | 1.000 ± 0.000 |
| Ligand | audio / flute | 571 | 1.000 ± 0.000 |
| Ligand | audio / pooled | 1 713 | 1.000 ± 0.000 |
| Ligand | MD baseline | 1 000 | 1.000 ± 0.000 |
| Ligand | Information-retention ratio | — | 1.000 |

Panel B. Leave-one-instrument-out generalisation
| Pair | Held-out instrument | n_train | n_test | Balanced accuracy |
| --- | --- | --- | --- | --- |
| Activation | piano | 1 130 | 565 | 0.594 |
| Activation | violin | 1 130 | 565 | 0.885 |
| Activation | flute | 1 130 | 565 | 0.750 |
| Activation | mean | — | — | 0.743 |
| Ligand | piano | 1 142 | 571 | 1.000 |
| Ligand | violin | 1 142 | 571 | 0.679 |
| Ligand | flute | 1 142 | 571 | 1.000 |
| Ligand | mean | — | — | 0.893 |

Panel C. Canonical correlation between MD and audio feature spaces
| Grouping | n_paired | r<sub>1</sub> | r<sub>2</sub> | r<sub>3</sub> | r<sub>4</sub> |
| --- | --- | --- | --- | --- | --- |
| Activation pair | 1 695 | 0.926 | 0.878 | 0.545 | 0.333 |
| Ligand pair | 1 713 | 0.995 | 0.883 | 0.652 | 0.591 |
| Three-state (supp.) | 2 409 | 0.994 | 0.855 | 0.825 | 0.561 |

**Figure 5.**
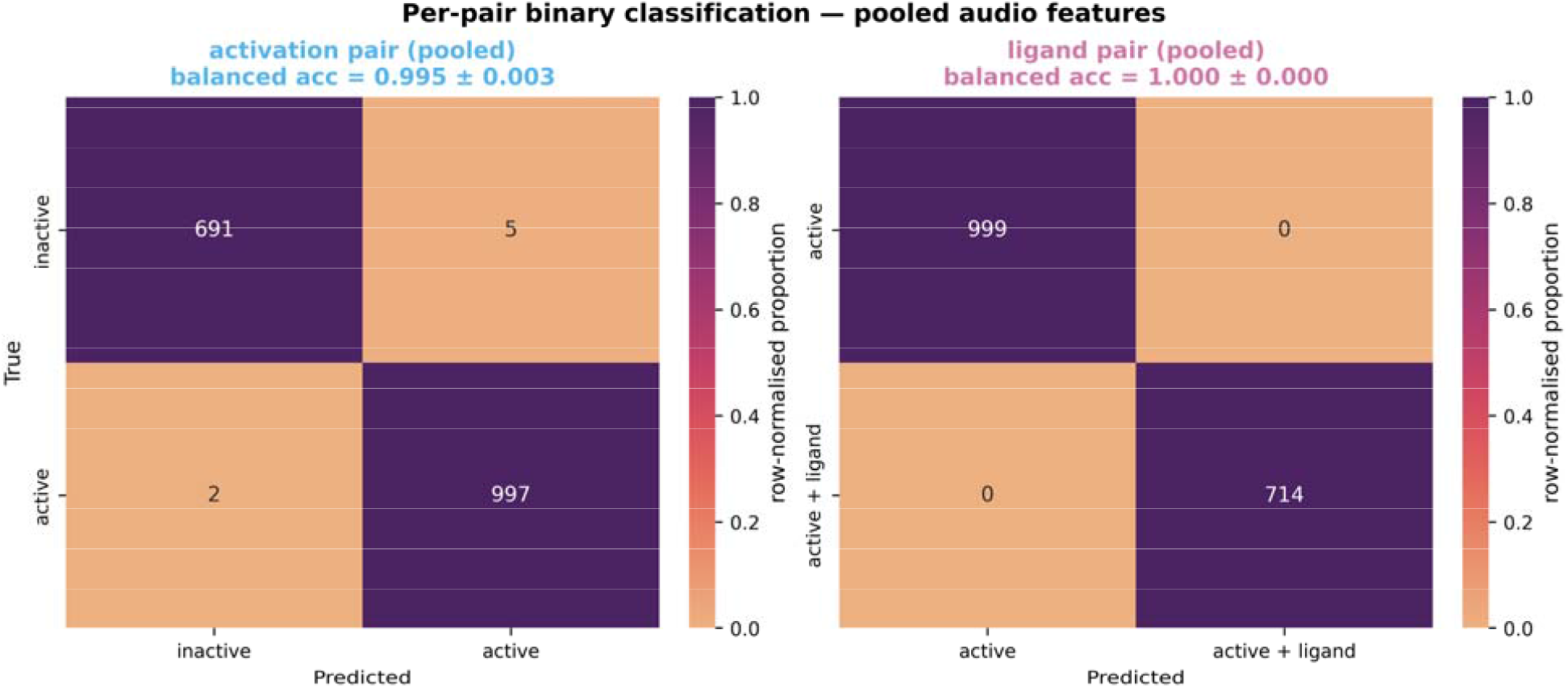
Per-pair binary classification — pooled audio features (headline result) Confusion matrices of the per-pair Random Forest classifiers trained on the windowed audio descriptor matrix (28 features per 1-s window, 0.5-s hop), pooled across all three instruments. (Left) Activation pair: classifier trained to distinguish inactive (*N* = 696 windows) from active apo (*N* = 999); 5-fold stratified cross-validated balanced accuracy 0.995 ± 0.003. (Right) Ligand pair: classifier trained to distinguish active apo (*N* = 999) from active + ligand (*N* = 714); balanced accuracy 1.000 ± 0.000 (perfect recovery). Cells annotated with raw counts; cell colour encodes row-normalised proportion (light → dark; sequential single-hue colour map). Panel titles are tinted with the corresponding pair colour (activation: sky blue; ligand: reddish purple).

To interpret these numbers we computed an information-retention ratio, the pooled audio classifier’s balanced accuracy divided by that of a Random Forest baseline trained directly on the MD features for the same pair (Methods 2.6). Per-pair MD baselines used only the MD features that carry information within each pair: four for the activation pair (TM3–TM6, NPxxY, DRY lock, Cα RMSD; ligand features are structurally undefined in apo systems and were dropped), and six for the ligand pair (the four above plus the two ligand-specific features, with biological sentinel values of contact_count = 0 and min_distance = 30 Å for the apo state). The MD-baseline classifiers reached 0.989 ± 0.006 (activation pair) and 1.000 ± 0.000 (ligand pair), yielding retention ratios of 1.006 for the activation pair and 1.000 for the ligand pair (Table 4).

These ratios are striking. For the ligand pair, the audio classifier exactly matches the upper bound set by direct access to MD features — the sonification preserves all of the MD’s binary state-discriminative information. For the activation pair, the audio classifier marginally exceeds the MD baseline; we attribute this to the fact that each 1-s audio window integrates approximately three to five MIDI notes (∼3–5 MD frames), allowing the classifier access to a short temporal context unavailable to the per-frame MD baseline. This is a property of the analysis window, not of the mapping per se; it does not imply the audio contains more information than the MD, but rather that the mapping followed by short-window aggregation produces a representation that is at least as discriminative as the raw frame-level MD vector. The 3-class supplementary analysis (Table S5) confirms the same conclusion in the joint problem (audio pooled balanced accuracy = 0.996 vs MD baseline 0.994). Together, these results provide a quantitative measure of information preservation in the biomolecular sonification mapping via cross-modal classification.

### 3.4. Cross-instrument generalisation is mixed: a faithful audit of timbre-independence

A faithful representation of MD-state should, in principle, generalise across the instruments used to render it: if the auditory signature truly carries MD information rather than timbre information, a classifier trained on (e.g.) piano + violin should recognise the state when tested on flute. We tested this by leave-one-instrument-out (LOIO) cross-validation per pair (Figure 7, Table S4).

**Figure 6.**
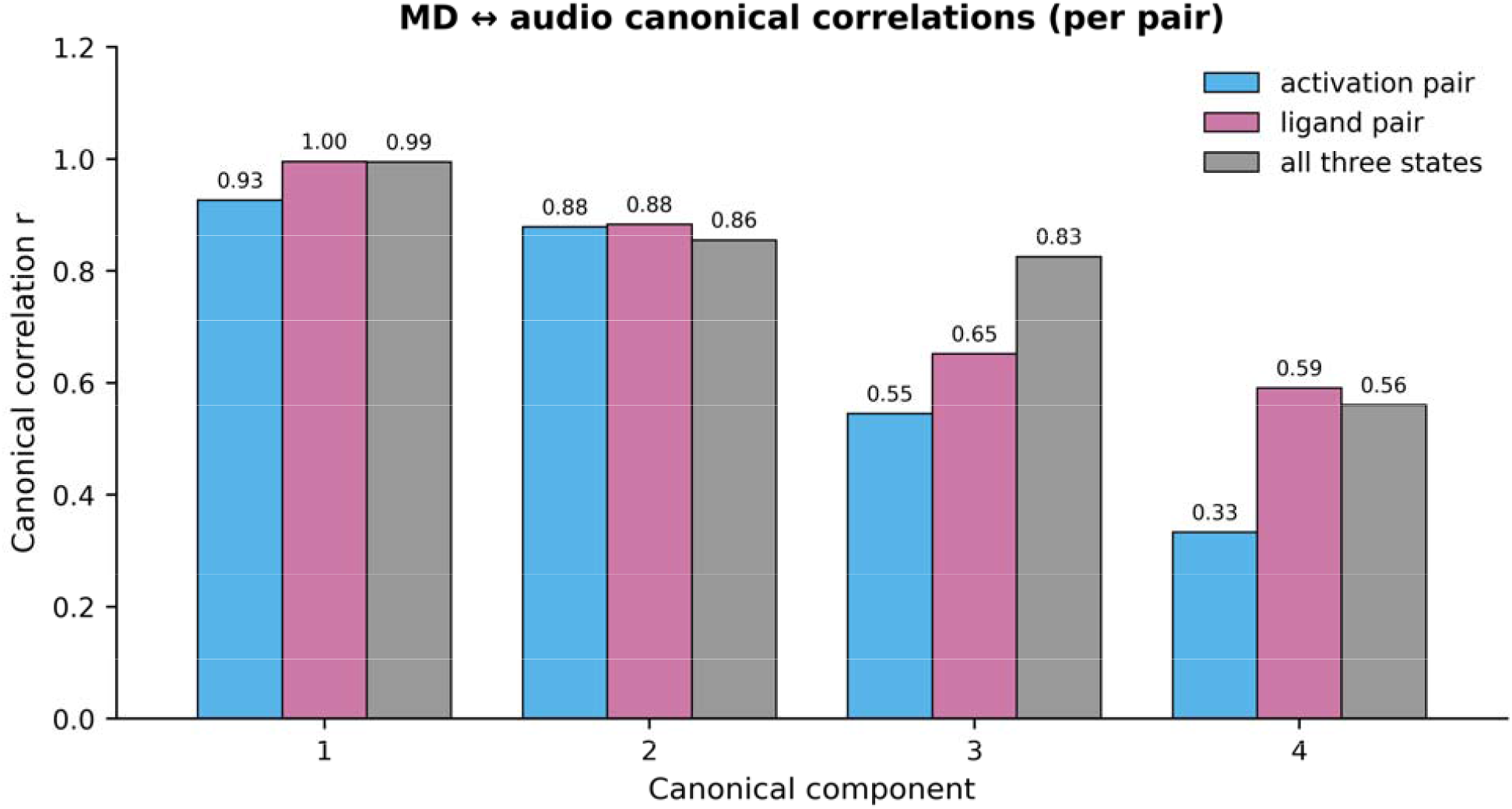
Canonical correlation between MD and audio feature spaces. First four canonical correlations *r*⍰ between standardised MD-feature vectors (per-pair valid subset: 4 features for the activation pair, 6 features for the ligand pair with biological sentinel imputation for apo ligand values) and standardised 28-D audio-descriptor vectors of the 1-s audio windows whose centre time aligns with each MD step. Bars grouped by canonical component (1–4) and coloured by pair: activation (sky blue, *n* = 1 695 paired samples), ligand (reddish purple, *n* = 1 713), and the supplementary three-state union (grey, *n* = 2 409). First canonical correlations reached r_1_ = 0.926 (activation), r_1_ = 0.995 (ligand), and r_1_ = 0.994 (three-state) — bilateral information-correspondence measure showing that, in the optimal projection of MD and audio feature spaces, the two representations share nearly all of their variance. Numeric values annotated above each bar. Canonical Correlation Analysis: sklearn.cross_decomposition.CCA, n_components = 4, max_iter 2000.

**Figure 7.**
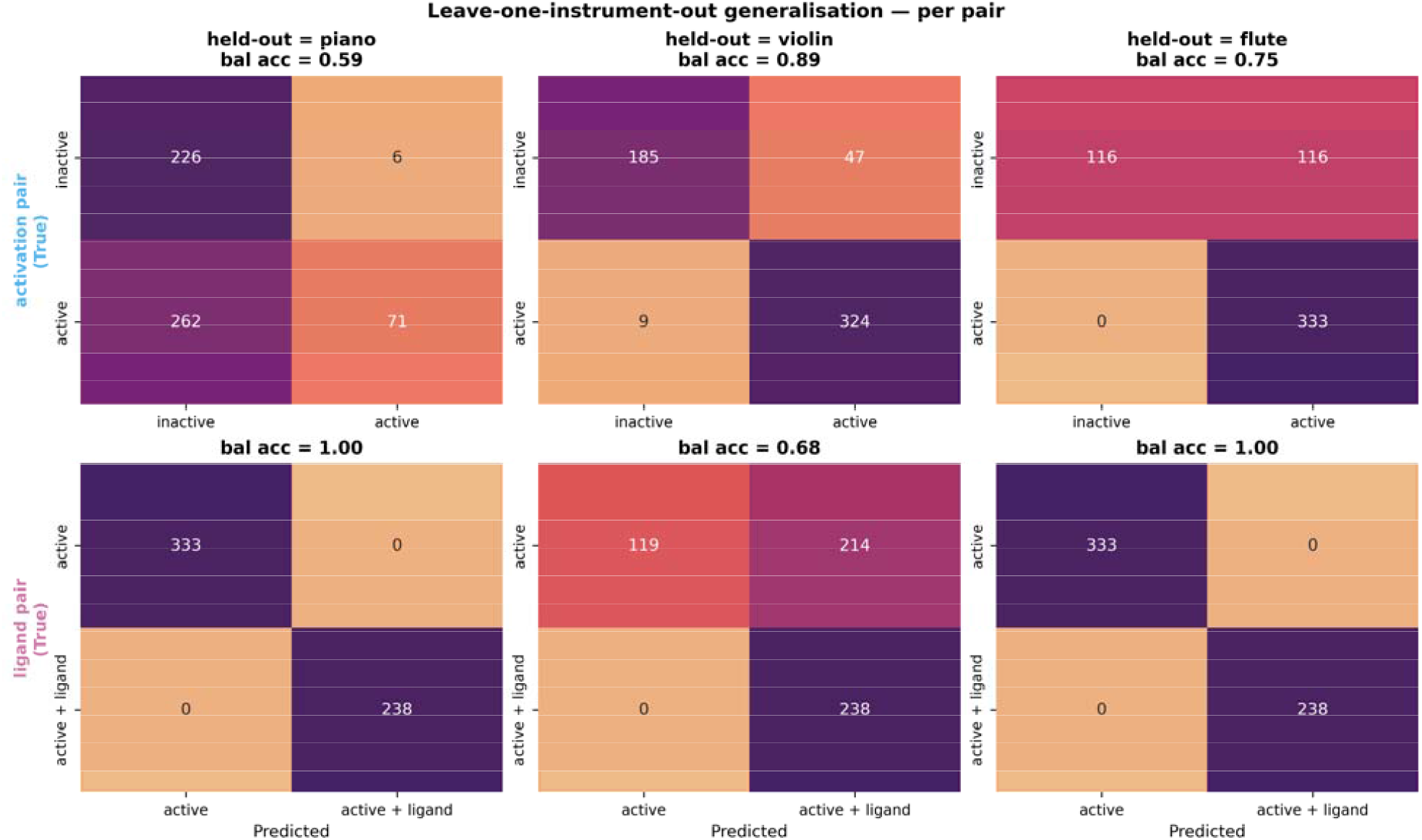
Leave-one-instrument-out (LOIO) generalisation per pair. Per-pair binary classification under leave-one-instrument-out cross-validation: a Random Forest classifier is trained on two of the three instruments’ windowed audio descriptors and tested on the held-out third. Rows: pair (top, activation; bottom, ligand). Columns: held-out instrument (piano, violin, flute). Each cell is a confusion matrix annotated with raw counts; cell colour encodes row-normalised proportion. Row labels are tinted with the corresponding pair colour. Within-instrument cross-validation balanced accuracies (Figure 5; Table 4 Panel A) are essentially perfect (0.988–1.000); under LOIO, balanced accuracy ranges from 0.594 to 1.000 with per-pair means 0.743 (activation) and 0.893 (ligand). The weakest cells — activation pair / held-out = piano and ligand pair / held-out = violin — show class-biased misclassification (e.g. 262/333 active piano windows assigned to “inactive”), characteristic of MFCC-derived audio descriptors whose timbre-specific manifolds do not fully overlap between training and held-out instruments.

The results are reported with their full nuance. Within instrument (the 5-fold scores in §3.3), generalisation is essentially perfect (0.988 – 1.000). Across instruments (LOIO), balanced accuracy ranges from 0.59 to 1.00. For the activation pair, LOIO yielded 0.594 (held-out = piano), 0.885 (held-out = violin), 0.750 (held-out = flute); for the ligand pair, 1.000 (held-out = piano), 0.679 (held-out = violin), 1.000 (held-out = flute). Per-pair mean LOIO balanced accuracies were 0.743 and 0.893 respectively — above the binary chance level of 0.500, but clearly below the within-instrument ceiling.

Inspection of the LOIO confusion matrices (Figure 7) clarifies the failure mode. The two weakest cases — activation pair / held-out = piano and ligand pair / held-out = violin — both exhibit systematic class-biased misclassification (e.g., the activation-pair model, having seen only violin and flute, assigns 262 of 333 active piano windows to “inactive”). This is the signature of an MFCC-based representation whose timbre-specific manifolds do not overlap between training and held-out instruments. The MD-state information is recoverable within each timbre — as evidenced by the 0.988 – 1.000 within-instrument scores — but its embedding in MFCC space carries an instrument-specific component that a classifier trained on two of three timbres cannot fully extrapolate.

This is, importantly, a limitation of the descriptor + classifier rather than of the sonification mapping. Three mitigation directions present themselves: (i) supplementing MFCCs with timbre-invariant descriptors (chroma, tonality, pitch-class profiles); (ii) per-instrument MFCC z-score normalisation before classifier training; and (iii) increasing the training-instrument panel (the present design uses only three GM patches). We do not implement these here so as to keep the framework’s primary computational claim — that the sonification mapping itself preserves MD-state information — cleanly separated from downstream representation-learning choices.

### 3.5. Canonical correlation analysis confirms bilateral information correspondence

Classification provides one notion of “the audio carries the MD information”; canonical correlation analysis (CCA) provides a complementary, classifier-free one. We aligned each 1-s audio window to the MD frame closest to its centre time (using the per-state MIDI-note start times reconstructed from the mapping table), yielding 1 695 paired (MD, audio) feature vectors for the activation pair, 1 713 for the ligand pair, and 2 409 for the full three-state union. We then fitted CCA between the (per-pair-valid) MD feature matrix and the 28-D audio descriptor matrix, recording the first four canonical correlations r⍰ (Figure 6, Table 4 Panel C).

The first canonical correlation reached r_1_ = 0.926 for the activation pair, r_1_ = 0.995 for the ligand pair, and r_1_ = 0.994 for the three-state union; second components followed at r_2_ ≈ 0.86 – 0.88 across all three groupings. These values quantify the strongest linear correspondence between the two representations: in the optimal projection of MD features and the optimal projection of audio descriptors, the two share nearly all of their variance. The slightly stronger r_1_ for the ligand pair (0.995 vs 0.926) is consistent with the larger absolute audio effect sizes observed in §3.2 — the percussion layer adds an audio mode that linearly tracks the ligand-contact MD feature with near-perfect fidelity.

Taken together with the classification results of §3.3, these correlations support a precise version of the central claim: the audio representation is not merely distinguishable between states (which would only require a single discriminative dimension), it linearly aligns with the MD representation in its dominant directions of variation. This is a strong information-correspondence measure for a deterministic mapping.

## 4. Discussion

### 4.1. Sonification as a complementary representation of molecular dynamics

The results presented in §3 provide a quantitative answer to a question that, to our knowledge, has not previously been asked of biomolecular sonification: does a deterministic mapping from MD features to musical parameters preserve the state-discriminative information present in the trajectories? For the β2-adrenergic receptor, the answer in this work is affirmative and quantitative — an audio-only Random Forest classifier reaches balanced accuracy 0.995 (activation pair) and 1.000 (ligand pair), recovering essentially all of the discriminative information available to a classifier trained directly on the MD features (retention ratios 1.006 and 1.000). A canonical correlation analysis supports the same conclusion in classifier-independent terms: the first canonical correlation between MD and audio feature spaces reaches r_1_ = 0.93 – 0.995, indicating that the two representations are closely aligned in their dominant directions of variation.

We frame this finding as a cross-modal representation result. The visualised MD trajectory and its rendered audio are two views of the same underlying conformational process, related by a deterministic mapping that — under the descriptor set tested here — preserves the relevant discriminative content. The auditory system parallel-processes temporal patterns that the visual system serialises through animation playback,^2^,^3^ and a single-pass audition of a 2-minute rendered trajectory presents the same conformational dynamics that a 500-frame cartoon animation would require an attentive viewer to interpret frame-by-frame. The sonification is not a replacement for structural visualisation but a complementary modality with different perceptual affordances: the snare-percussion layer that fires only in the ligand-bound state, for example, communicates an interaction event whose visual analogue would require a researcher to manually count receptor–ligand contacts across hundreds of frames.

### 4.2. A biological observation that emerged from the sonification design

A specific observation arising from the per-feature analysis warrants attention. The NPxxY motif of the seventh transmembrane helix shows the largest single-feature effect size in both pair contrasts (|Cliff’s δ| = 0.76 for the activation pair, 0.75 for the ligand pair), but the direction of the ligand-pair effect is opposite to the other features. While TM3–TM6 distance, DRY ionic-lock distance, and Cα RMSD all continue to increase from active apo to active + ligand (consistent with deeper engagement of the G-protein-binding pocket), NPxxY RMSD decreases from 4.08 ± 1.45 Å to 2.08 ± 0.68 Å upon agonist binding. The same feature is therefore the slowest in the active apo audio (longest notes) and the most stable in the ligand-bound audio (consistent, faster note durations).

This is consistent with the established model in which orthosteric agonists rigidify NPxxY into a G-protein-binding-competent rotamer.^23,24,17,16^ The mapping rendered this rigidification audibly: a listener attending to note duration alone can distinguish the active apo regime (slow, exploratory) from the active + ligand regime (fast, locked). Sonification did not discover this finding — it was already in the per-frame feature table — but the auditory representation provides a complementary channel through which the same observation becomes perceptually evident.

### 4.3. Limitations and methodological caveats

We report three limitations honestly. First, our design uses a single MD trajectory per state (the GPCRMD reference trajectories), and the statistical tests of §3.1 – §3.2 therefore quantify variability within a single MD run rather than across independent replicas. Multiple-replica MD with the present pipeline would distinguish stochastic from state-specific contributions to the audio signature; the per-state mapping table format produced by Notebook 02 supports such an extension without changes to the audio rendering or classifier code.

Second, cross-instrument generalisation is mixed. The within-instrument classifiers reach 0.988 – 1.000, but LOIO balanced accuracy ranges from 0.594 to 1.000 (per-pair means 0.74 and 0.89). The failure mode (§3.4) is class-biased confusion characteristic of an MFCC manifold that does not extrapolate cleanly to an unseen timbre. We deliberately did not introduce timbre-invariant descriptors (chroma, pitch-class profiles) or per-instrument normalisation because doing so would conflate the present claim — that the mapping itself is information-preserving — with downstream representation-learning choices. A separate study comparing MFCC, chroma, and learnt audio embeddings under LOIO would address this directly.

Third, the sonification mapping is heuristic, not optimised. We selected biologically interpretable axis assignments (activation distance → pitch, motif dynamics → tempo, ionic-lock → harmonic intensity, contacts → percussion), but no algorithmic criterion was used to choose them. The framework’s information-retention ratio (§3.3) and canonical correlations (§3.5) provide a post hoc scoring system that could, in future work, be used as the optimisation target of a search over mapping choices, turning mapping design into a measurable rather than aesthetic problem.

Finally, this is a computational study of representation fidelity, not of human perception. The question of whether human listeners can discriminate the rendered audio signatures above chance — and how this discrimination scales with musical training, listening time, and stimulus length — is the natural complementary study, but lies outside the scope of the present computational claim.

### 4.4. Applications and outlook

The pipeline is immediately applicable to any GPCR for which a multi-state MD ensemble exists; transferring it requires only updating the residue-range selection strings to the new sequence numbering. Extension beyond GPCRs is similarly direct for any allosteric system in which a small set of activation-related geometric features can be defined (kinases, ion channels, riboswitches, allosteric metabolic enzymes). Three application contexts seem especially well-suited to the framework presented here.

Exploratory pattern discovery on long trajectories. Microsecond-to-millisecond MD trajectories are now routinely produced^10,11^ but remain difficult to scan visually. A real-time audio rendering of a long trajectory provides a parallel perceptual stream in which transient events (lock break, allosteric rearrangement, ligand approach) can be flagged for closer visual inspection. The percussion-only-when-bound design used here for β2AR generalises to any threshold event in MD.

Accessibility. Blind and low-vision researchers have, to date, had no primary-modality access to MD trajectories beyond textual summaries. A sonification with documented information-retention guarantees provides such access. The HTML player accompanying this work (Notebook 04) bundles audio, video, and synchronised playback so the same artefact serves sighted and non-sighted users.

Education and outreach. Receptor activation as a phenomenon is conceptually subtle for introductory students; the audible difference between the inactive, active, and active + ligand renders provides an additional handle that complements the visual cartoon. The Colab pipeline and pre-rendered audio files are released openly to support educational reuse.

Beyond these application contexts, the framework offers a general methodological proposal: information-retention ratio and canonical correlation between source-feature and rendered-audio-descriptor spaces can serve as standardised, interpretation-free scoring criteria for sonification mappings, applicable to any deterministic mapping in any scientific domain.

## 5. Conclusions

We have presented a computationally validated sonification framework for GPCR molecular dynamics, applied as a proof of concept to the three GPCRMD β2-adrenergic receptor reference trajectories. The framework itself is system-agnostic and transfers to any GPCR (or any allosteric MD system) by adjusting only the residue-selection strings. Under a single deterministic mapping rule, the activation-related geometric features extracted from β2AR MD are translated into three perceptually distinct audio signatures whose state-of-origin can be recovered, by a Random Forest classifier trained on audio features alone, at balanced accuracy ≥ 0.995 within each instrument and with information-retention ratios of 1.006 and 1.000 relative to the MD-feature baselines for the two designed pair contrasts. Canonical correlation between MD and audio feature spaces reaches r_1_ = 0.93 – 0.995, providing classifier-free evidence that the mapping yields an auditory representation closely aligned with the underlying MD dynamics. Cross-instrument generalisation under leave-one-instrument-out testing is mixed (mean balanced accuracy 0.74 – 0.89), reflecting timbre-specific MFCC manifolds rather than loss of MD signal; this is reported transparently and identifies a clear methodological direction for follow-up work on timbre-invariant audio descriptors.

The biological analysis embedded in the pipeline also reproduces, in an audibly perceptible form, the established stabilisation of the NPxxY motif upon orthosteric agonist binding, supplying a concrete example of a known structural finding rendered into a complementary perceptual channel. The framework — pipeline, audio assets, classification + canonical-correlation evaluation, and synchronised HTML player — is released openly to support reproduction, extension to other GPCRs and allosteric systems, and use in accessibility and educational contexts. Beyond the specific β2AR application, we propose information-retention ratio and canonical correlation between source-feature and rendered-audio descriptor spaces as standardised, interpretation-free scoring criteria for any deterministic sonification mapping in scientific data analysis.

## Author Contributions

E.Y.: Conceptualization, Methodology, Software, Formal analysis, Investigation, Writing Original Draft, Writing Review & Editing.

## Notes

The author declares no competing financial interest.

## Acknowledgments

We thank the GPCRMD consortium for providing the β2AR reference trajectories used in this work.

## Data and Code Availability

All MD trajectory data used in this work is publicly available from the GPCRMD repository (https://gpcrmd.org) under Dynamic IDs 11, 116, and 117. All processed data (per-frame features, MIDI scores, audio renders, classification outputs) and all analysis code (five Jupyter / Colab notebooks) are openly available at Zenodo 10.5281/zenodo.20343439 and mirrored at GitHub https://github.com/eygpcr/gpcr-sonification-beta2ar. A synchronised HTML player and pre-rendered audio + video assets are included in the Zenodo deposit.

## Supporting Information

A separate Supporting Information document contains six supplementary tables (Tables S1 – S6: per-state MIDI parameter summary; PCA loadings; full per-pair MWU on all 28 audio descriptors; leave-one-instrument-out confusion matrices; 3-class supplementary classification; per-pair MD-feature baseline classifier details) and ten supplementary figures (Figures S1 – S10: QC plots; diagnostic MIDI pitch contour; MD feature time series; spectrograms for violin and flute timbres; ligand-pair cross-instrument grid; piano-roll comparisons; audio-feature PCA; 3-state MD-feature PCA; 3-class pooled confusion matrix).

## References

(1) Kramer, G., Ed. Auditory Display: Sonification, Audification, and Auditory Interfaces; Addison-Wesley: Reading, MA, 1994.

(2) Hermann, T.; Hunt, A.; Neuhoff, J. G., Eds. The Sonification Handbook; Logos Verlag: Berlin, 2011.

(3) Walker, B. N.; Nees, M. A. Theory of Sonification. In The Sonification Handbook; Hermann, T., Hunt, A., Neuhoff, J. G., Eds.; Logos Verlag: Berlin, 2011; pp 9–39.

(4) Cooke, J.; Díaz-Merced, W.; Foran, G.; Hannam, J.; García, B. Exploring Data Sonification to Enable, Enhance, and Accelerate the Analysis of Big, Noisy, and Multi-Dimensional Data. Proc. Int. Astron. Union 2019, 14, 251–256.

(5) Díaz-Merced, W. L. Sound for the Exploration of Space Physics Data. PhD Thesis, University of Glasgow, 2013.

(6) Hermann, T. Taxonomy and Definitions for Sonification and Auditory Display. Proc. 14th Int. Conf. Auditory Display (ICAD 2008), Paris, France, 2008; pp 1–8.

(7) Yu, C.-H.; Qin, Z.; Martin-Martinez, F. J.; Buehler, M. J. A Self-Consistent Sonification Method to Translate Amino Acid Sequences into Musical Compositions and Application in Protein Design Using Artificial Intelligence. ACS Nano 2019, 13 (7), 7471–7482.

(8) Franjou, S. L.; Milazzo, M.; Yu, C.-H.; Buehler, M. J. Sounds Interesting: Can Sonification Help Us Design New Proteins? Expert Rev. Proteomics 2019, 16 (11–12), 875–879.

(9) Larsen, P.; Gilbert, J. Microbial Bebop: Creating Music from Complex Dynamics in Microbial Ecology. PLoS ONE 2013, 8 (3), e58119.

(10) Shaw, D. E.; Maragakis, P.; Lindorff-Larsen, K.; Piana, S.; Dror, R. O.; Eastwood, M. P.; Bank, J. A.; Jumper, J. M.; Salmon, J. K.; Shan, Y.; Wriggers, W. Atomic-Level Characterization of the Structural Dynamics of Proteins. Science 2010, 330 (6002), 341–346.

(11) Lindorff-Larsen, K.; Piana, S.; Dror, R. O.; Shaw, D. E. How Fast-Folding Proteins Fold. Science 2011, 334 (6055), 517–520.

(12) Dunn, J.; Clark, M. A. Life Music: The Sonification of Proteins. Leonardo 1999, 32 (1), 25–32.

(13) Bywater, R. P.; Middleton, J. N. Melody Discrimination and Protein Fold Classification. Heliyon 2016, 2 (10), e00175.

(14) Garcia-Ruiz, M. A.; Gutierrez-Pulido, J. R. An Overview of Auditory Display to Assist Comprehension of Molecular Information. Interact. Comput. 2006, 18 (4), 853–868.

(15) Hauser, A. S.; Attwood, M. M.; Rask-Andersen, M.; Schiöth, H. B.; Gloriam, D. E. Trends in GPCR Drug Discovery: New Agents, Targets and Indications. Nat. Rev. Drug Discov. 2017, 16 (12), 829–842.

(16) Rasmussen, S. G. F.; DeVree, B. T.; Zou, Y.; Kruse, A. C.; Chung, K. Y.; et al. Crystal Structure of the β2 Adrenergic Receptor–Gs Protein Complex. Nature 2011, 477 (7366), 549–555.

(17) Manglik, A.; Kim, T. H.; Masureel, M.; Altenbach, C.; Yang, Z.; Hilger, D.; et al. Structural Insights into the Dynamic Process of β2-Adrenergic Receptor Signaling. Cell 2015, 161 (5), 1101–1111.

(18) Liu, X.; Masoudi, A.; Kahsai, A. W.; Huang, L.-Y.; Pani, B.; et al. Mechanism of β2AR Regulation by an Intracellular Positive Allosteric Modulator. Science 2019, 364 (6447), 1283–1287.

(19) Rodríguez-Espigares, I.; Torrens-Fontanals, M.; Tiemann, J. K. S.; et al. GPCRmd Uncovers the Dynamics of the 3D-GPCRome. Nat. Methods 2020, 17 (8), 777–787.

(20) Kobilka, B. K. G Protein Coupled Receptor Structure and Activation. Biochim. Biophys. Acta 2007, 1768 (4), 794–807.

(21) Lefkowitz, R. J. A Brief History of G-Protein Coupled Receptors (Nobel Lecture). Angew. Chem. Int. Ed. 2013, 52 (25), 6366–6378.

(22) Rosenbaum, D. M.; Rasmussen, S. G. F.; Kobilka, B. K. The Structure and Function of G-Protein-Coupled Receptors. Nature 2009, 459 (7245), 356–363.

(23) Latorraca, N. R.; Venkatakrishnan, A. J.; Dror, R. O. GPCR Dynamics: Structures in Motion. Chem. Rev. 2017, 117 (1), 139–155.

(24) Dror, R. O.; Arlow, D. H.; Maragakis, P.; Mildorf, T. J.; Pan, A. C.; Xu, H.; Borhani, D. W.; Shaw, D. E. Activation Mechanism of the β2-Adrenergic Receptor. Proc. Natl. Acad. Sci. U.S.A. 2011, 108 (46), 18684–18689.

(25) Michaud-Agrawal, N.; Denning, E. J.; Woolf, T. B.; Beckstein, O. MDAnalysis: A Toolkit for the Analysis of Molecular Dynamics Simulations. J. Comput. Chem. 2011, 32 (10), 2319–2327.

(26) Gowers, R. J.; Linke, M.; Barnoud, J.; et al. MDAnalysis: A Python Package for the Rapid Analysis of Molecular Dynamics Simulations. Proc. 15th Python in Science Conf. (SciPy 2016) 2016, 98–105.

(27) Raffel, C.; Ellis, D. P. W. Intuitive Analysis, Creation and Manipulation of MIDI Data with pretty_midi. 15th Int. Soc. for Music Information Retrieval Conf. Late-Breaking Demo, 2014.

(28) McFee, B.; Raffel, C.; Liang, D.; Ellis, D. P. W.; McVicar, M.; Battenberg, E.; Nieto, O. librosa: Audio and Music Signal Analysis in Python. Proc. 14th Python in Science Conf. 2015, 18–25.

(29) Romano, J.; Kromrey, J. D.; Coraggio, J.; Skowronek, J. Appropriate Statistics for Ordinal Level Data: Should We Really Be Using t-Test and Cohen’s d for Evaluating Group Differences on the NSSE and Other Surveys? Annu. Meeting Florida Assoc. of Institutional Research, 2006; pp 1–33.

(30) Benjamini, Y.; Hochberg, Y. Controlling the False Discovery Rate: A Practical and Powerful Approach to Multiple Testing. J. R. Stat. Soc. B 1995, 57 (1), 289–300.

(31) Brodersen, K. H.; Ong, C. S.; Stephan, K. E.; Buhmann, J. M. The Balanced Accuracy and Its Posterior Distribution. Proc. 20th Int. Conf. Pattern Recognition (ICPR) 2010, 3121–3124.

