## Supplementary for "A Sonification Framework for GPCR Molecular Dynamics: Auditory Signatures of β2-Adrenergic Receptor"

Ekrem Yasar

### Contents of this Supporting Information

- Table S1. Per-state MIDI parameter summary
- Table S2. PCA loadings for the four common MD features
- Table S3. Acoustic feature Mann–Whitney U tests — all 28 descriptors × 2 pairs
- Table S4. Leave-one-instrument-out confusion matrices — per-pair binary
- Table S5. Three-class supplementary classification
- Table S6. Per-pair MD-feature baseline classifier — extended
- Figures S1 – S10. Quality-control plots, time-series, spectrograms, PCA, and 3-class confusion matrix (file list below)

### Table S1. Per-state MIDI parameter summary

*Statistics over the 500 sonification steps per state. MIDI note range C3 – C6 (48 – 84); note duration linearly scaled from NPxxY-RMSD; velocity scaled from Cα RMSD. State order follows the activation continuum.*

| State | n | Pitch (MIDI note) | Duration (s) | Velocity (MIDI) |
| --- | --- | --- | --- | --- |
| Inactive | 500 | 57.7 ± 4.3 [48 – 67] | 0.214 ± 0.038 [0.12 – 0.35] | 72.0 ± 7.3 [40 – 90] |
| Active (apo) | 500 | 64.0 ± 5.1 [55 – 79] | 0.315 ± 0.069 [0.12 – 0.45] | 77.9 ± 8.5 [40 – 89] |
| Active + ligand | 500 | 71.7 ± 6.4 [50 – 84] | 0.219 ± 0.033 [0.12 – 0.28] | 84.8 ± 10.7 [40 – 110] |

*Each cell: mean ± SD [min – max]. Pitch values shifted upward with activation; note duration peaks in the active apo state (NPxxY-RMSD maximum) and shortens in the ligand-bound state (NPxxY-RMSD stabilisation); velocity increases monotonically with activation.*

### Table S2. PCA loadings for the four common MD features

*Principal components fitted on standardised MD features pooled across all three states (Notebook 03, §11). Two components retained.*

| MD feature | PC1 | PC2 |
| --- | --- | --- |
| TM3–TM6 intracellular distance | +0.649 | +0.218 |
| NPxxY motif RMSD | −0.376 | +0.563 |
| DRY ionic-lock distance | +0.652 | +0.233 |
| Cα RMSD | −0.107 | +0.762 |

*PC1 is dominated by the two intracellular helix-opening distances (TM3–TM6 and DRY lock) — the canonical activation axis. PC2 loads primarily on Cα RMSD (global flexibility) and NPxxY motif RMSD, separating the active apo ensemble (high motif dynamics) from the ligand-bound ensemble (stabilised motif).*

### Table S3. Acoustic feature Mann–Whitney U tests — all 28 descriptors × 2 pairs

*Per-pair MWU on every windowed audio descriptor (1-s windows, 0.5-s hop, ~1 700 windows per pair pooled across the three instruments). Cliff's δ as effect size; Benjamini–Hochberg-corrected q-values. Rows sorted by |Cliff's δ| (descending). Significant rows at q < 0.05 are marked *.*

#### Panel A. Activation pair (inactive vs active)

| Audio descriptor | mean_A | mean_B | U | q (BH) | \|δ\| | sig |
| --- | --- | --- | --- | --- | --- | --- |
| rms_mean | 0.026 | 0.036 | 61 119 | 4.7 × 10⁻¹⁸³ | 0.824 | * |
| mfcc12_std | 4.62 | 6.16 | 144 543 | 8.8 × 10⁻⁹³ | 0.584 | * |
| mfcc8_std | 4.68 | 6.17 | 149 837 | 4.3 × 10⁻⁸⁸ | 0.569 | * |
| mfcc12_mean | −3.63 | −8.08 | 530 708 | 1.2 × 10⁻⁷⁵ | 0.527 | * |
| mfcc13_std | 4.56 | 5.75 | 166 457 | 3.6 × 10⁻⁷⁴ | 0.521 | * |
| mfcc7_std | 4.87 | 6.07 | 185 384 | 8.5 × 10⁻⁶⁰ | 0.467 | * |
| mfcc6_std | 5.47 | 6.88 | 193 173 | 2.4 × 10⁻⁵⁴ | 0.444 | * |
| mfcc10_std | 4.97 | 6.10 | 196 645 | 5.3 × 10⁻⁵² | 0.434 | * |
| mfcc9_std | 4.84 | 5.92 | 197 795 | 3.0 × 10⁻⁵¹ | 0.431 | * |
| mfcc6_mean | 6.67 | 1.06 | 495 250 | 9.0 × 10⁻⁵⁰ | 0.425 | * |
| zcr_std | 7.0 × 10⁻³ | 9.0 × 10⁻³ | 200 614 | 2.0 × 10⁻⁴⁹ | 0.423 | * |
| mfcc10_mean | −2.82 | −6.65 | 492 158 | 8.6 × 10⁻⁴⁸ | 0.416 | * |
| mfcc9_mean | 1.01 | −2.30 | 487 642 | 5.9 × 10⁻⁴⁵ | 0.403 | * |
| mfcc4_std | 8.65 | 10.09 | 216 416 | 1.0 × 10⁻³⁹ | 0.377 | * |
| mfcc11_std | 4.64 | 5.46 | 218 044 | 9.0 × 10⁻³⁹ | 0.373 | * |
| mfcc11_mean | −3.96 | −6.78 | 472 467 | 4.5 × 10⁻³⁶ | 0.359 | * |
| rms_std | 5.2 × 10⁻³ | 6.6 × 10⁻³ | 223 971 | 1.9 × 10⁻³⁵ | 0.356 | * |
| mfcc5_mean | 21.49 | 17.28 | 451 540 | 1.9 × 10⁻²⁵ | 0.299 | * |
| mfcc5_std | 6.64 | 7.61 | 258 220 | 3.0 × 10⁻¹⁹ | 0.257 | * |
| mfcc8_mean | −2.68 | −4.79 | 433 731 | 6.2 × 10⁻¹⁸ | 0.248 | * |
| mfcc7_mean | 5.46 | 0.64 | 430 636 | 8.9 × 10⁻¹⁷ | 0.239 | * |
| zcr_mean | 0.024 | 0.028 | 268 138 | 1.6 × 10⁻¹⁵ | 0.229 | * |
| mfcc13_mean | −7.28 | −8.75 | 425 967 | 4.2 × 10⁻¹⁵ | 0.225 | * |
| mfcc1_mean | −429.60 | −399.27 | 280 005 | 1.3 × 10⁻¹¹ | 0.195 | * |
| mfcc1_std | 25.42 | 27.58 | 287 901 | 2.3 × 10⁻⁹ | 0.172 | * |
| mfcc3_std | 13.10 | 14.30 | 290 100 | 8.7 × 10⁻⁹ | 0.166 | * |
| mfcc4_mean | 31.52 | 37.09 | 290 279 | 9.5 × 10⁻⁹ | 0.165 | * |
| mfcc2_mean | 211.40 | 205.32 | 404 715 | 1.1 × 10⁻⁸ | 0.164 | * |
| spec_rolloff_std | 458.71 | 500.26 | 315 588 | 1.5 × 10⁻³ | 0.092 | * |
| spec_bandwidth_mean | 1558.30 | 1612.22 | 319 671 | 5.4 × 10⁻³ | 0.080 | * |
| spec_rolloff_mean | 2174.35 | 2248.97 | 319 706 | 5.4 × 10⁻³ | 0.080 | * |
| spec_centroid_std | 242.00 | 251.98 | 321 946 | 1.1 × 10⁻² | 0.074 | * |
| mfcc3_mean | −25.70 | −31.29 | 325 220 | 2.6 × 10⁻² | 0.065 | * |
| mfcc2_std | 12.66 | 12.93 | 326 127 | 3.2 × 10⁻² | 0.062 | * |
| spec_centroid_mean | 1055.19 | 1101.74 | 330 205 | 8.2 × 10⁻² | 0.050 | ns |
| spec_bandwidth_std | 394.14 | 386.69 | 351 717 | 6.8 × 10⁻¹ | 0.012 | ns |

#### Panel B. Ligand pair (active apo vs active + ligand)

| Audio descriptor | mean_A | mean_B | U | q (BH) | \|δ\| | sig |
| --- | --- | --- | --- | --- | --- | --- |
| mfcc1_mean | −399.27 | −279.40 | 0 | 9.5 × 10⁻²⁷² | 1.000 | * |
| mfcc5_mean | 17.28 | −14.69 | 713 175 | 9.5 × 10⁻²⁷² | 1.000 | * |
| mfcc2_mean | 205.32 | 158.72 | 710 829 | 2.3 × 10⁻²⁶⁸ | 0.993 | * |
| spec_rolloff_std | 500.26 | 1370.29 | 2 932 | 7.8 × 10⁻²⁶⁸ | 0.992 | * |
| mfcc1_std | 27.58 | 58.04 | 2 955 | 7.8 × 10⁻²⁶⁸ | 0.992 | * |
| spec_rolloff_mean | 2248.97 | 4369.76 | 4 132 | 3.8 × 10⁻²⁶⁶ | 0.988 | * |
| rms_mean | 0.036 | 0.062 | 4 393 | 8.1 × 10⁻²⁶⁶ | 0.988 | * |
| spec_centroid_std | 251.98 | 543.53 | 8 456 | 8.4 × 10⁻²⁶⁰ | 0.976 | * |
| rms_std | 6.6 × 10⁻³ | 1.7 × 10⁻² | 8 638 | 1.4 × 10⁻²⁵⁹ | 0.976 | * |
| spec_centroid_mean | 1101.74 | 1959.77 | 9 109 | 6.2 × 10⁻²⁵⁹ | 0.974 | * |
| spec_bandwidth_mean | 1612.22 | 2509.61 | 9 854 | 7.2 × 10⁻²⁵⁸ | 0.972 | * |
| mfcc13_mean | −8.75 | 2.70 | 23 232 | 1.7 × 10⁻²³⁸ | 0.935 | * |
| mfcc8_mean | −4.79 | −14.42 | 669 528 | 3.2 × 10⁻²¹⁰ | 0.877 | * |
| mfcc5_std | 7.61 | 11.98 | 55 957 | 2.7 × 10⁻¹⁹⁴ | 0.843 | * |
| mfcc4_std | 10.09 | 16.05 | 57 226 | 1.1 × 10⁻¹⁹² | 0.840 | * |
| zcr_std | 9.0 × 10⁻³ | 1.4 × 10⁻² | 84 718 | 3.1 × 10⁻¹⁵⁹ | 0.762 | * |
| mfcc6_std | 6.88 | 9.81 | 90 351 | 8.8 × 10⁻¹⁵³ | 0.747 | * |
| mfcc9_mean | −2.30 | −7.81 | 570 913 | 2.0 × 10⁻⁹⁹ | 0.601 | * |
| mfcc10_mean | −6.65 | −12.17 | 567 546 | 2.2 × 10⁻⁹⁶ | 0.591 | * |
| zcr_mean | 0.028 | 0.035 | 146 112 | 4.4 × 10⁻⁹⁶ | 0.590 | * |
| mfcc11_std | 5.46 | 6.58 | 187 499 | 1.4 × 10⁻⁶² | 0.474 | * |
| mfcc4_mean | 37.09 | 56.08 | 221 543 | 1.5 × 10⁻⁴⁰ | 0.379 | * |
| mfcc8_std | 6.17 | 6.79 | 242 383 | 1.9 × 10⁻²⁹ | 0.320 | * |
| mfcc6_mean | 1.06 | 4.57 | 256 621 | 6.5 × 10⁻²³ | 0.280 | * |
| mfcc2_std | 12.93 | 11.70 | 449 384 | 6.7 × 10⁻²⁰ | 0.260 | * |
| mfcc12_mean | −8.08 | −6.04 | 281 399 | 1.3 × 10⁻¹³ | 0.211 | * |
| mfcc12_std | 6.16 | 5.85 | 421 104 | 2.4 × 10⁻¹⁰ | 0.181 | * |
| spec_bandwidth_std | 386.69 | 421.96 | 303 271 | 1.6 × 10⁻⁷ | 0.150 | * |
| mfcc10_std | 6.10 | 6.56 | 303 917 | 2.2 × 10⁻⁷ | 0.148 | * |
| mfcc3_mean | −31.29 | −44.02 | 402 944 | 5.6 × 10⁻⁶ | 0.130 | * |
| mfcc7_std | 6.07 | 5.79 | 389 764 | 1.3 × 10⁻³ | 0.093 | * |
| mfcc7_mean | 0.64 | −3.71 | 387 597 | 2.6 × 10⁻³ | 0.087 | * |
| mfcc3_std | 14.30 | 13.85 | 386 948 | 3.1 × 10⁻³ | 0.085 | * |
| mfcc11_mean | −6.78 | −6.23 | 333 969 | 2.6 × 10⁻² | 0.064 | * |
| mfcc13_std | 5.75 | 5.72 | 368 228 | 2.6 × 10⁻¹ | 0.032 | ns |
| mfcc9_std | 5.92 | 5.83 | 365 146 | 4.0 × 10⁻¹ | 0.024 | ns |

*Total significant descriptors: activation pair 34 / 36, ligand pair 34 / 36 (q < 0.05 BH). The ligand pair shows 17 descriptors with |Cliff's δ| > 0.7 (effectively perfect separation), driven by the snare-percussion track that is triggered only when the ligand-contact count is non-zero.*

### Table S4. Leave-one-instrument-out confusion matrices — per-pair binary

*Expanded form of Table 4 Panel B. Random Forest trained on two of three instruments, tested on the held-out third. Rows are the true class; columns are the predicted class. Class-biased misclassifications (a single column dominating in the off-diagonal) reflect non-overlap of the held-out instrument's MFCC manifold with the training instruments.*

#### Activation pair

| Held-out | Pred → inactive (True: inact. / active) | Pred → active (True: inact. / active) | Balanced acc. |
| --- | --- | --- | --- |
| piano | 226 / 262 | 6 / 71 | 0.594 |
| violin | 185 / 9 | 47 / 324 | 0.885 |
| flute | 116 / 0 | 116 / 333 | 0.750 |

#### Ligand pair

| Held-out | Pred → active (True: active / active+lig.) | Pred → active+lig. (True: active / active+lig.) | Balanced acc. |
| --- | --- | --- | --- |
| piano | 333 / 0 | 0 / 238 | 1.000 |
| violin | 119 / 0 | 214 / 238 | 0.679 |
| flute | 333 / 0 | 0 / 238 | 1.000 |

*Bold counts in the off-diagonals (e.g. 262, 214, 116) highlight the class-bias direction. The other classifiers correctly assign every window.*

### Table S5. Three-class supplementary classification

*Same Random Forest framework as Table 4 Panel A, but trained as a single 3-class classifier (chance = 0.333). Reported per instrument and pooled. Included for completeness; shows that the framework generalises beyond the two designed pair contrasts.*

#### Panel A. Within-instrument 5-fold CV

| Model | n_samples | Balanced accuracy |
| --- | --- | --- |
| 3-class / piano | 803 | 0.992 ± 0.008 |
| 3-class / violin | 803 | 0.997 ± 0.006 |
| 3-class / flute | 803 | 0.998 ± 0.003 |
| 3-class / pooled | 2 409 | 0.996 ± 0.002 |
| MD baseline / 3-class | 1 500 | 0.994 ± 0.005 |

#### Panel B. 3-class LOIO

| Held-out instrument | n_train | n_test | Balanced accuracy |
| --- | --- | --- | --- |
| piano | 1 606 | 803 | 0.728 |
| violin | 1 606 | 803 | 0.726 |
| flute | 1 606 | 803 | 0.805 |
| mean | — | — | 0.753 |

*The 3-class pooled audio classifier (0.996) reaches the MD-feature baseline (0.994), confirming the binary per-pair conclusion at the joint multi-class level. LOIO exhibits the same instrument-specific MFCC-manifold limitation as in the binary case.*

### Table S6. Per-pair MD-feature baseline classifier — extended

*The MD-feature baseline classifier used to compute the information-retention ratio in Table 4. Per-pair valid features only (apo states with structurally-undefined ligand features dropped for the activation pair, biological-sentinel imputation for the ligand pair).*

| Model | n_samples | Features used | Balanced accuracy |
| --- | --- | --- | --- |
| MD baseline / activation | 1 000 | RMSD_Ca, TM3_TM6_distance, NPxxY_RMSD, DRY_ionic_lock (4) | 0.989 ± 0.006 |
| MD baseline / ligand | 1 000 | RMSD_Ca, TM3_TM6_distance, NPxxY_RMSD, DRY_ionic_lock, Ligand_min_distance*, Ligand_contact_count* (6) | 1.000 ± 0.000 |
| MD baseline / 3-class | 1 500 | All 6 MD features* | 0.994 ± 0.005 |

** For ligand-pair and 3-class models, the apo state's structurally-undefined ligand features are imputed with biologically motivated sentinels: Ligand_contact_count = 0 (literal zero contacts in apo) and Ligand_min_distance = 30 Å ("no-ligand" sentinel, much greater than any physical bound distance).*

### Supplementary Figures

**Figure S1.** QC plots from Notebook 01 — per-feature time series across all three states (6 panels: Cα RMSD, TM3–TM6 distance, NPxxY RMSD, DRY ionic lock, ligand minimum distance, ligand contact count). Files: QC_*.png

**Figure S2.** Diagnostic MIDI pitch contour per pair (Notebook 02). File: diag_midi_note_sequence_per_pair.png

**Figure S3.** MD feature time series per pair (Notebook 03). Files: fig_pair_activation_features_timeseries.png, fig_pair_ligand_features_timeseries.png

**Figure S4.** Activation pair: side-by-side spectrograms for violin and flute. Files: fig_pair_activation_spectrograms_violin.png, fig_pair_activation_spectrograms_flute.png

**Figure S5.** Ligand pair: side-by-side spectrograms for violin and flute. Files: fig_pair_ligand_spectrograms_violin.png, fig_pair_ligand_spectrograms_flute.png

**Figure S6.** Ligand pair: cross-instrument robustness grid (2 × 3, piano / violin / flute × active / active + ligand). File: fig_pair_ligand_cross_instrument_grid.png

**Figure S7.** Piano-roll comparisons per pair. Files: fig_pair_activation_piano_roll.png, fig_pair_ligand_piano_roll.png

**Figure S8.** PCA of windowed audio feature space (per-state colour, per-instrument marker shape). File: fig_audio_features_pca.png

**Figure S9.** 3-state MD-feature PCA. File: fig_feature_pca.png

**Figure S10.** 3-class pooled confusion matrix (supplementary analysis). File: fig_classification_confusion_3class_pooled.png
